# Learning the differences: a transfer-learning approach to predict antigen immunogenicity and T-cell receptor specificity

**DOI:** 10.1101/2022.12.06.519259

**Authors:** Barbara Bravi, Andrea Di Gioacchino, Jorge Fernandez-de-Cossio-Diaz, Aleksandra M. Walczak, Thierry Mora, Simona Cocco, Rémi Monasson

## Abstract

Antigen immunogenicity and the specificity of binding of T-cell receptors to antigens are key properties underlying effective immune responses. Here we propose diffRBM, an approach based on transfer learning and Restricted Boltzmann Machines, to build sequence-based predictive models of these properties. DiffRBM is designed to learn the distinctive patterns in amino acid composition that, one the one hand, underlie the antigen’s probability of triggering a response, and on the other hand the T-cell receptor’s ability to bind to a given antigen. We show that the patterns learnt by diffRBM allow us to predict putative contact sites of the antigen-receptor complex. We also discriminate immunogenic and non-immunogenic antigens, antigen-specific and generic receptors, reaching performances that compare favorably to existing sequence-based predictors of antigen immunogenicity and T-cell receptor specificity. More broadly, diffRBM provides a general framework to detect, interpret and leverage selected features in biological data.

## 1 Introduction

T cells play an essential role in the immune response to pathogens and malignancies. Killer T cells are activated following the binding of their surface receptors (T-cell receptors or TCRs) to short portions of pathogen-related proteins (peptide antigens) that are presented by class I major histocompatibility complexes (MHCs) forming the peptide-MHC epitope (pMHC).

Only a fraction of peptides presented by the MHC are immunogenic, meaning that they possess biochemical properties that can promote a T-cell response [Sette et al., 1994]. Accurate prediction of immunogenicity is crucial to the successful identification of microbial antigens and cancer neoantigens (antigens carrying cancer-related mutations) that help develop vaccines and immune-based cancer therapies. A very recent systematic assessment [Buckley et al., 2022] of the available models to identify immunogenic targets from pathogens and cancers reports suboptimal overall performances, with none of the models able to substantially improve beyond pure MHC-presentation prediction when evaluated on immunogenic peptides from a new virus (SARS-CoV-2). The largest-scale validation [Wells et al., 2020] of existing computational pipelines for neoantigen discovery has highlighted a general lack of consensus among their predictions and a rather low average success rate, with only 6% of the predicted neoantigens validated as truly immunogenic.

We also know that a given pMHC epitope elicits the response of only specific small subsets of the human T-cell repertoire. Predicting the molecular composition of TCRs that have the potential to be reactive to a given epitope is a difficult computational problem that is yet not fully solved despite numerous recent advances [Gielis et al., 2019, Jokinen et al., 2019, Springer et al., 2020, Montemurro et al., 2021, Weber et al., 2021]. Improvements in predicting immunogenicity and TCR specificity would have direct consequences for medical applications, including the study of an individual’s infection history from their T-cell repertoire and personalized adoptive T-cell therapy for cancer treatment.

Both antigen immunogenicity and epitope-specificity of T-cell receptors have a molecular-level component. They result from specific physico-chemical constraints on the sequence composition of antigens and T-cell receptors. Immunogenic antigens and epitope-specific receptors display an enrichment in specific patterns of amino acid composition. For example, several works have shown enrichment in hydrophobic [Chowell et al., 2015, Riley et al., 2019] and aromatic [Schmidt et al., 2021] residues in immunogenic peptides, compared to all presented peptides (which are predominantly non-binders of TCRs). TCRs specifically responding to the same peptide are also characterized by convergent amino acid motifs [Cinelli et al., 2017, Dash et al., 2017, Glanville et al., 2017, Pogorelyy et al., 2019, Huang et al., 2020, Shomuradova et al., 2020, Mayer-Blackwell et al., 2021, Minervina et al., 2021, Goncharov et al., 2022], whose retrieval is the focus of several clustering approaches [Dash et al., 2017, Glanville et al., 2017, Meysman et al., 2019, Pogorelyy and Shugay, 2019, Thakkar and Bailey-Kellogg, 2019, Mayer-Blackwell et al., 2021, Valkiers et al., 2021]. These patterns are observed in addition to the sequence patterns that are more broadly shared across antigens and TCRs. These general patterns reflect baseline constraints ensuring viability and function (ensuring that TCRs are structurally stable and have the basic binding properties allowing them to pass thymic selection, or ensuring that antigens have high binding affinity to the presenting HLA protein). An outstanding question is how to disentangle sequence pattern enrichment underlying immunogenicity and TCR epitope specificity from the stronger statistical signatures stemming from these baseline constraints. This separation could generate insight into the molecular basis of antigen immunogenicity and epitope specificity and could enable predicting them, at least to some extent, from the sequence alone.

To tackle this question, we here introduce a strategy of ‘differential learning’ within the unsupervised learning scheme of Restricted Boltzmann Machines [Hinton, 2002, Hinton and Salakhutdinov, 2006], which we call diffRBM (differential Restricted Boltzmann Machine). DiffRBM relies on a transfer learning procedure, where we first learn general background distributions of antigen or TCR sequences, and then capture, from few sequence data, the distinctive features that confer them, respectively, immunogenicity or epitope specificity. We next interpret the sequence patterns learnt by diffRBM, and assess its performance compared to existing computational tools.

## 2 A differential learning approach: diffRBM

In this paper, we apply diffRBM to build models of peptide immunogenicity and T-cell binding to specific peptides. The modeling setting to which diffRBM is applicable is more general. It could be applied to any data that have some distinctive features compared to a much larger pool of data endowed with the baseline properties. We will refer to these two different sets of data as ‘selected’ and ‘background’ datasets (Figure 1A-B). In our example, they correspond to sets of immunogenic and only presented antigens in the case of the model of immunogenicity, or antigen-specific and bulk-repertoire TCRs in the case of the model of TCR epitope specificity (Figure 1C-D). Scenarios where it is useful to predict distinctive properties of a subset of data that are important for their functional selection against a background are numerous in biology, with the increasing deployment of SELEX [Ellington and Szostak, 1990, Tuerk and Gold, 1990, Sola et al., 2020] and directed evolution protocols [Jäckel et al., 2008, Packer and Liu, 2015, Arnold, 2018].

**Figure 1:**
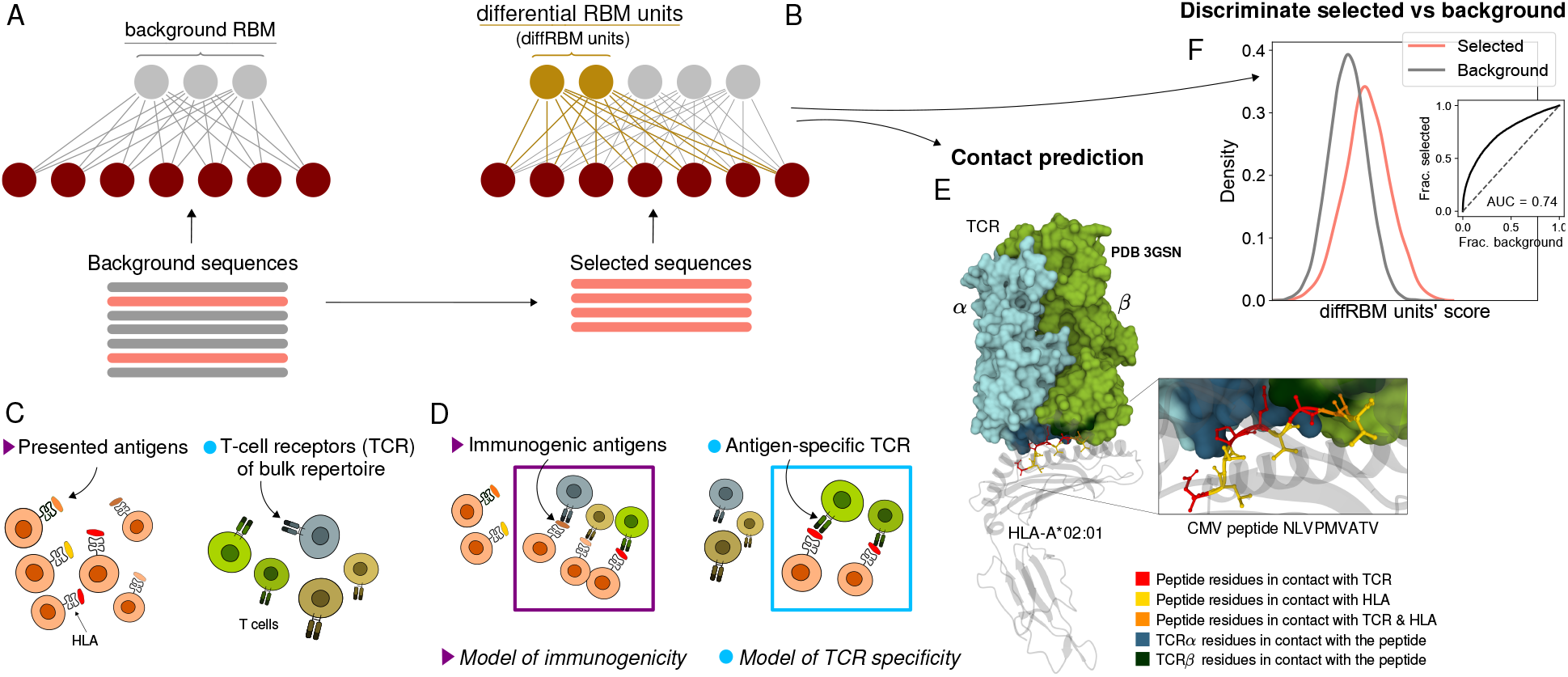
Cartoon of the differential RBM (diffRBM) learning approach. A: The parameters of background RBM (gray) are learnt from the ‘background’ sequence dataset. B: The diffRBM units (gold) are learnt from a small subset of ‘selected’ sequences. C: We consider the application of diffRBM to modeling peptide immunogenicity or T-cell receptor (TCR) antigen specificity, whereby the background dataset consists, respectively, of all antigens presented by a given Human Leukocyte Antigen class I complex (HLA) or of generic TCRs from the bulk repertoire. D: The selected sequences correspond to HLA-specific antigens validated to be immunogenic or to TCRs that are antigen-specific responders. The inferred parameters associated to the diffRBM units allow one to identify putative contact positions in the peptide-HLA-TCR structure (E) and more generally to assign scores that distinguish the selected from the background sequences (F). E is an example of a peptide-HLA-TCR structure for the CMV peptide NLVPMVATV (PDB-ID:3GSN), where the contact points along the peptide and the TCR are highlighted in different colors (image obtained with Mol* [Sehnal et al., 2018]).

Our differential approach is akin to the machine learning technique known as transfer learning, whereby a model learnt for one task is transferred to the second task in such a way that the information embedded in the first model facilitates the learning of the second model. Transfer learning has been recently applied in the context of immune receptors, to model antigen-specific T-cell receptors [Isacchini et al., 2021, Lu et al., 2021] and antibodies [Akbar et al., 2022], but not in the context of modeling antigen immunogenicity. The benefit of a transfer learning approach is two-fold. First, it allows one to leverage the background data availability to control for overfitting. Using the background RBM, which is pre-trained on the larger background dataset as a starting point for learning the diffRBM units, reduces the amount of parameters and data features to infer from the selected dataset (typically much smaller than the background dataset). A second advantage, particularly relevant to the interpretable modeling of biological data, is the fact that the diffRBM units focus on the differences relative to the background, disentangling in this way the data features that make them a selected subset with distinctive properties. As a result, scores based on the diffRBM units can distinguish selected data from the background data (Figure 1F). In the context of modeling antigen immunogenicity and TCR specificity, the diffRBM units and their parameters can also identify antigen-TCR contact points in the three-dimensional molecular structure (Figure 1E).

## 3 Results

### 3.1 DiffRBM model of antigen immunogenicity

Several approaches have been proposed to predict antigen immunogenicity [Calis et al., 2013, Trolle and Nielsen, 2014, Chowell et al., 2015, Ogishi and Yotsuyanagi, 2019, Gao et al., 2020, Schmidt et al., 2021, Lin et al., 2021, Buckley et al., 2022], some of them specifically developed for computational pipelines of neoantigen discovery [Łuksza et al., 2017, Smith et al., 2019, Riley et al., 2019, Schenck et al., 2019, Schaap-Johansen et al., 2021]. The model [Calis et al., 2013] is currently the most used resource for MHC class I immunogenicity prediction, being both implemented by the IEDB tool (http://tools.iedb.org/immunogenicity/) for immunogenicity predictions and integrated in the T cell-antigen interaction prediction by the NetTepi server [Trolle and Nielsen, 2014]. It relies on the simple assumption of position-independence of the main biophysical properties underlying immunogenicity. Some of these approaches are based on a preliminary choice of peptide positions [Schmidt et al., 2021] or properties (such as hydrophobicity [Chowell et al., 2015]), known or assumed to be important for recognition by TCRs. The diffRBM approach we propose is a sequence-based method that does not require an *ad hoc* selection of residue positions or properties and does not rely on the assumption of peptide position independence.

We collected from the Immune Epitope Database (IEDB) [Vita et al., 2019] sets of peptides that elicited a T-cell reaction in T cell assays (referred to as ‘immunogenic’) and sets of peptides that were not T-cell-reactive (‘non-immunogenic’). The peptide-presenting MHCs are specialized proteins coded by highly polymorphic human genes called Human Leukocyte Antigen (HLA) gene. We selected only peptides presented by 3 HLA-I alleles (HLA-A*02:01, HLA-B*35:01, HLA-B*07:02). We chose these HLA-I alleles since they are associated to at least 200 immunogenic peptides in IEDB and at least one TCR-pMHC structure in the Protein Data Bank (Methods). We trained diffRBM for each set of HLA-restricted immunogenic peptides in two steps, by training first a background RBM on samples of antigens presented by one specific HLA via the RBM-based algorithm RBM-MHC [Bravi et al., 2021b], and next by training the diffRBM units on the set of immunogenic antigens of the same HLA type (Figures 2A, S2). Background RBM can predict scores of presentation on the specific HLA under consideration, while the diffRBM units predict scores of immunogenicity.

**Figure 2:**
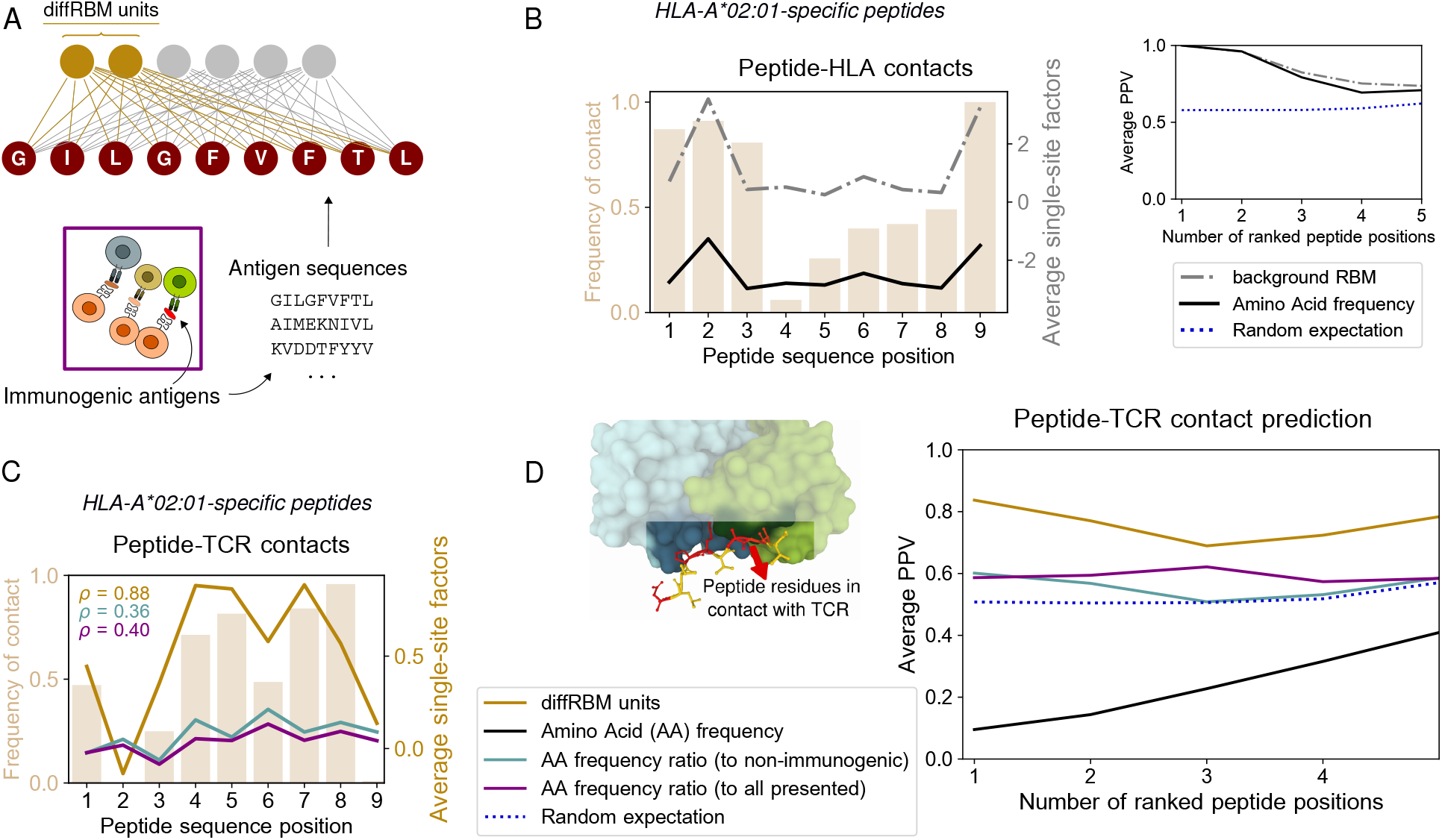
DiffRBM model of immunogenicity and structural interpretation of its parameters. A: The diffRBM units are learnt from HLA-specific peptides annotated as immunogenic. B: From the background dataset (HLA-A*02:01-presented antigens), we estimate the expected amino acid frequency at each position; here we plot its average value for the 41 HLA-A*02:01-restricted peptides in the available crystal structures (black line) and we compare it to the magnitude of the single-site factors from background RBM (HLA-A*02:01-specific presentation model), averaged over the 41 peptides. The trend of the amino acid frequency and single-site factors resembles the one of the frequency of contacts with the HLA across the 41 structures for each peptide position. Right inset: Average Positive Predictive Value (PPV) for the prediction of peptide positions in contact with the HLA as a function of the number of ranked positions. The PPV is averaged over the 41 HLA-A*02:01-restricted peptides. The random expectation (blue dotted line) gives the average PPV over a uniformly random prediction of each set of contacts (Methods). C: Same as B, where peptide contact positions are the ones with the TCR and the single-site factors are estimated from either the diffRBM units modeling immunogenicity or amino acid frequency ratios (ratio of position-specific amino acid frequency in immunogenic peptides to non-immunogenic peptides or all presented peptides, see legend in D). *ρ* denotes the coefficient of correlation between the contact frequency distribution and the magnitude of single-site factors. Contact positions with the HLA (respectively TCR) are defined as the peptide positions whose distance to the residues of the HLA (respectively TCR) in the resolved structure is within 3.5 (respectively 4) Å. D: PPV for the prediction of peptide positions in contact with the TCR as a function of the peptide positions ranked by the models’ single-site factors. For each peptide, predictions are made using the corresponding HLA-specific model of immunogenicity. The average is over 46 structures (4 for HLA-B*35:01, 41 for HLA-A*02:01, 1 for HLA-B*07:02). All average PPVs are reweighed by a term accounting for the sequence similarity between peptide entries (Methods, Figures S4A-B, S5A-B).

Differently from some existing approaches to modeling immunogenicity [Calis et al., 2013, Chowell et al., 2015, Schmidt et al., 2021], we train *HLA-specific* models. Preliminary inspection of the datasets revealed that patterns of amino acid enrichment differ between immunogenic peptides presented by different HLAs, apart from some general trend in terms of dominant amino acid properties [Calis et al., 2013, Chowell et al., 2015, Riley et al., 2019, Schmidt et al., 2021, Łuksza et al., 2022]. This is true also when we restrict to the peptide positions known to be relevant for immunogenicity [Schmidt et al., 2021, Rudolph et al., 2006, Calis et al., 2012, Schmidt et al., 2021, Łuksza et al., 2022] (Figure 3A). For instance, the extent to which enrichment in hydrophobicity can discriminate immunogenic from non-immunogenic peptides was observed to vary across HLAs [Buckley et al., 2022], since presentation by certain HLAs alone might be associated to hydrophobic residues, supporting HLA-specific strategies to model immunogenicity. On the practical side, cross-HLA imbalances in the training sets, due to the substantial variation in the availability of immunogenic peptides across HLAs, were found to skew predictions toward the most characterized HLAs, in particular toward HLA-A*02:01 [Buckley et al., 2022].

**Figure 3:**
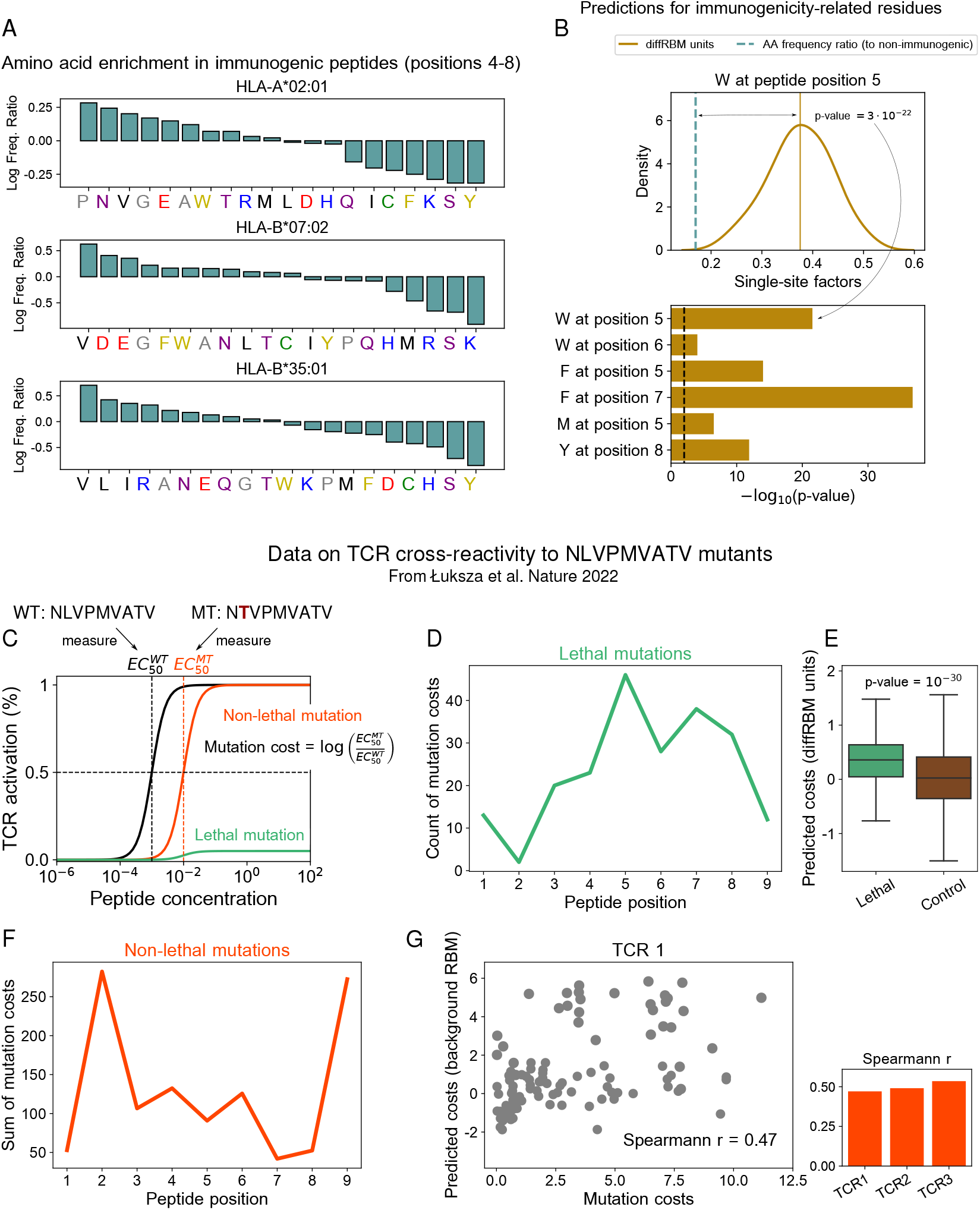
DiffRBM units encode molecular-level features of immunogenicity. A: Enrichment in amino acid usage in immunogenic peptides relative to non-immunogenic ones calculated as a log frequency ratio across the central positions (4-8) for each HLA type. The color code used in the x-axis labels indicates amino acid properties: red = negatively charged (E, D), blue = positively charged (H, K, R), purple = non-charged polar hydrophilic (N, T, S, Q), golden = aromatic (F, W, Y), black = aliphatic hydrophobic (I, L, M, V), green = cysteine (C), grey = tiny (A, G, P) or other. B: Distribution of the diffRBM single-site factors evaluated across the HLA-A*02:01-specific immunogenic sequences with W at position 5. For all amino acid/position combinations that were found to be key to immunogenicity, the diffRBM single-site factors assume predominantly positive values (see also Figure S6A). This means that diffRBM predicts a positive contribution to immunogenicity of these key residues, in agreement with observations. Taking into account the sequence context of these residues is essential here: the single-site factors given by the immunogenic *vs* non-immunogenic amino acid frequency ratio, which do not include the sequence context (Methods), predict a much lower contribution to immunogenicity, as indicated by the p-values of their difference with respect to the average of the diffRBM single-site factors distribution. The p-value is estimated from the one-sample T-test and the black dashed line marks the reference p-value=0.01. C: Cartoon illustration of the TCR activation curves obtained in [Łuksza et al., 2022] for the wild-type (*WT*) peptide NLVPMVATV and its single-site mutants (*MT*). 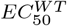 and 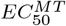 indicate the half-point peptide concentration of the TCR activation curve for respectively *WT* and *MT* (Methods). D: The total count of lethal mutations’ costs (214 out of the 513 TCR-mutant combinations) is plotted per mutated peptide position. E: The distribution of predictions of the cost of lethal mutations by diffRBM units is significantly shifted towards positive values (Methods). F: Sum of the costs of non-lethal mutations (299 out of the 514 TCR-mutant combinations) per mutated peptide position. G: Experimental mutation costs *vs* the mutation costs predicted by background RBM for the non-lethal mutations in the context of one TCR (TCR1). The Spearmann correlation coefficient *r* is comparable across the 3 TCRs, with a p-value ≤ 10^−6^ for all of them (Figure S6B).

Equipped with these single-alleles models of immunogenicity, we can perform two tasks, the prediction of peptide sites in contact with the TCR (Figure 1E) and the classification of immunogenic peptides against non-immunogenic ones (Figure 1F).

### 3.2 Validation of model predictions against TCR-pMHC structures

Since the diffRBM units are inferred to capture the distinctive patterns of peptide immunogenicity, we hypothesized that the associated inferred parameters can be informative about the actual structural properties of the pMHC-TCR complex. To test this hypothesis, we collected a set of resolved crystal structures publicly available in the Protein Data Bank [Berman et al., 2000] describing peptides in complex with the presenting HLA molecule and a cognate TCR. For each of these TCR-pMHC complexes, we estimated the peptide positions in contact with the TCR and the HLA (Methods, Table S1). Figures 2B-C show the frequency of contacts at each peptide position with the HLA (B) and the TCR (C) in peptides presented by HLA-A*02:01 (the HLA allele to which the large majority of structures is available, 41 over 46 structures, see Methods). These contact frequency distributions highlight that positions 2 and 9 (and 1 and 3 to a lower extent) are the anchor sites for the binding of the peptide to the HLA-I protein (Figure 2B), while central positions (4-8) tend to be in contact to the TCR (Figure 2C), consistently with the analyses of structures reported in [Rudolph et al., 2006, Calis et al., 2012, Schmidt et al., 2021, Milighetti et al., 2021]. Previous measures of TCR functional avidity with mutant peptides indicate that amino acid changes at the peptide central positions impact the most T-cell activation [Hoof et al., 2010, Schmidt et al., 2021, Łuksza et al., 2022], suggesting that these positions are important for TCR response.

#### Anchor sites of peptide-HLA binding can be inferred from the amino acid statistics

We first focus on HLA-peptide binding. Anchor sites for the bond with the HLA constrain the amino acid usage at those positions across peptides presented by that same HLA, increasing the frequency of the amino acids required for binding (*e.g*., I, L, V at positions 2 and 9 of HLA-A*02:01 ligands). As a result, the sites occupied by the high-frequency amino acids at those positions are typically anchor sites (Figure 2B). We can translate this simple observation into a recipe to predict peptide contacts with the HLA. For each peptide sequence from the TCR-pMHC-A*02:01 structures, we ranked sequence positions by the frequency of the amino acids estimated from the background dataset of HLA-A*02:01-presented peptides. We took the top ranking positions as putative anchor sites and we verified this prediction against the true contacts by calculating an average Positive Predictive Value (PPV), see the inset of Figure 2B and Methods. This average PPV gives the fraction of ranked positions that correspond to the true contacts averaged over HLA-A*02:01-specific peptides from the available structures as a function of the number of ranked positions (*i.e*. predicted contacts). The PPV is equal to its maximal value, 1, for the first ranked position and stays close to it for the successive ones (Figure 2B inset, Figure S5).

We can also formulate a model-based prediction of contact sites. We took the background RBM parameters, which are learnt to reproduce the amino acid statistics in HLA-A*02:01-presented peptides, and we used them to define peptide site-specific quantities that we call ‘single-site factors’. These correspond essentially to the background RBM log probabilities of single sites conditional on the rest of the sequence (Methods). Large single-site factors detect the peptide-HLA binding anchor sites, similarly to the amino acid frequency (Figure 2B), hence anchor sites can be predicted as the top ranking sequence positions according to the single-site factors. The average PPV of this prediction is comparable to the amino acid frequency-based prediction (Figure 2B inset).

#### The diffRBM single-site factors flag up peptide positions important for immunogenicity

Also in sets of immunogenic peptides we expect the statistics at the contact positions with the TCR to reflect the constraint of being in contact, when compared to the statistics of all presented peptides. These constraints are captured by the parameters linked to the diffRBM units, as they are learnt in such a way as to pick up the statistical differences with respect to all presented peptides. In analogy to the prediction of HLA-peptide binding via background RBM, we introduce what we call ‘diffRBM single-site factors’ to predict the single residue potential to establish a contact with the TCR. The diffRBM single-site factors are defined from the parameters of the diffRBM units in such a way as to give approximately log odds-ratios between the full RBM and the background RBM probabilities of a certain residue conditional on the rest of the sequence (equation 17 in Methods). Once evaluated on a peptide, these model-dependent terms provide a measure of the predicted contribution to immunogenicity of the amino acid at each peptide position. Since they are based on the diffRBM units, which are connected simultaneously to all sequence sites, the diffRBM single-site factors deliver effectively single-site predictions while accounting for the sequence context given by all the other sites (hence, for the same position and amino acid, their values can vary between sequences).

Figure 2C shows the average magnitude of the diffRBM single-site factors evaluated on the 41 peptides binding to HLA-A*A02:01, which identifies positions 4-8 as the most relevant for immunogenicity. The pattern of positional contact frequency with the TCR in the same figure supports the structural interpretation of the model’s prediction in terms of binding between peptide and TCR.

The diffRBM units’ prediction recovers the pattern of positions important for immunogenicity without restricting *a priori* the input sequences to a subset of peptide positions already known or assumed to be involved in TCR binding. This is in contrast with existing approaches [Calis et al., 2013, Schmidt et al., 2021] that take advantage of considerations on peptide-HLA binding to choose *a priori* what positions to retain in the formulation of the immunogenicity model. [Calis et al., 2013] excludes the positions that are most commonly the anchor sites in the binding to the HLA (1,2 and 9). [Schmidt et al., 2021] learns a classifier of immunogenicity using only the HLA-specific set of positions (for example 5-8 for HLA-A*02:01) estimated to have a minimal impact on HLA affinity based on the statistical information content of HLA-I binding motifs.

#### The diffRBM single-site factors predict peptide contact positions with the TCR

We ranked sequence positions by the single-site factors’ magnitude for each peptide in the TCR-pMHC complexes, using the HLA-specific model corresponding to the peptide’s HLA type, and we took the highest ranking positions as predicted contact points. The peptide-averaged PPV for this prediction as a function of the ranked positions (Figure 2D) indicates a model’s predictive power substantially higher than the random expectation (Methods).

Differential amino acid usage between immunogenic and non-immunogenic peptides at single positions, each considered independently of each other, has been essential to inspect what biochemical properties contribute to immunogenicity [Calis et al., 2013, Chowell et al., 2015] and has been leveraged to build predictors of immunogenicity [Calis et al., 2013]. We compared the prediction by the diffRBM units to such predictions based on differential amino acid usage (Figure 2C-D). Here the single-site factors are given by the log ratio between the position-specific amino acid frequency in immunogenic peptides of a given HLA type and the one in the set of either all presented peptides or the non-immunogenic peptides with the same HLA type (Methods). Thus these single-site factors describe, under an independent-site assumption, the enrichment in amino acid usage in immunogenic peptides relative to all presented peptides or non-immunogenic ones respectively. The diffRBM units outperform these predictions based on amino acid frequency ratios, as quantified by the average PPV (Figures 2D, S4). For the HLA-A*02:01 peptides, we also found that the magnitude of the diffRBM single-site factors across positions correlates with the pattern of contact frequency better than predictions based on amino acid frequency ratios (Figure 2C).

Finally, Figure 2D shows that a prediction based only on the frequency of an amino acid in the dataset of immunogenic peptides, similarly to the dataset of all presented peptides, tends to identify as putative contacts the anchor sites of binding to the HLA, leading to completely incorrect predictions concerning the binding sites to the TCR.

### 3.3 DiffRBM encodes molecular-level features of immunogenicity

Next we assessed whether our model predictions of the residues’ contribution to immunogenicity, based on single-site factors, are consistent with previous findings [Piepenbrink et al., 2013, Chowell et al., 2015, Riley et al., 2019, Schmidt et al., 2021, Łuksza et al., 2022]. Inspecting the amino acid enrichment with respect to non-immunogenic antigens at the common peptide-TCR contact points (positions 4-8) reveals similarities but also differences compared to other accounts of immunogenicity (Figure 3A). Some of the trends visible in Figure 3A resemble the ones found in previous studies, for instance: the bias, in immunogenic sequences, towards hydrophobic amino acids [Chowell et al., 2015, Riley et al., 2019, Łuksza et al., 2022, Buckley et al., 2022] (especially valine, denoted by symbol V); the abundance of glutamic acid (E) [Calis et al., 2013] and, to a more moderate extent, of tryptophan (W) [Schmidt et al., 2021, Łuksza et al., 2022]; the depletion of the small polar amino acid serine (S) and the positively charged amino acid lysine (K), consistently with the observations respectively in [Chowell et al., 2015] and [Schmidt et al., 2021]. We noted also a number of discrepancies, presumably due to the use of datasets of immunogenic and non-immunogenic peptides that were updated with respect to previously available ones and restricted to sequences of only three selected HLA types. The most striking is the under-representation of the aromatic amino acids phenylalanine (F) and tyrosine (Y), particularly severe in HLA-A*02:01, in contrast with the experiment-based observations in [Schmidt et al., 2021, Piepenbrink et al., 2013]. (Note however that also other analyses performed on IEDB data, like [Calis et al., 2013, Chowell et al., 2015], did not flag up a significant enrichment in Y).

#### The diffRBM single-site factors recover the positive contribution to immunogenicity of key residues

We then verified whether these apparent discrepancies in terms of site-specific amino acid abundances in our IEDB-derived dataset persisted at the level of model predictions. We considered a few combinations of amino acids and positions along the peptide that were suggested to play a crucial role in T-cell reactivity and binding (in the context of HLA-A*02:01 epitopes) based on structural or functional analyses. For instance, W at position 6 and F at position 7 [Schmidt et al., 2021], as well as Y at position 8 [Piepenbrink et al., 2013], were observed to form a variety of stabilizing interactions with the TCR. Testing functional avidity of TCRs against peptides harbouring single-point mutations, [Schmidt et al., 2021] detected that F and W at position 5 triggered the strongest activation signal, while [Łuksza et al., 2022] found that the substitution of methionine (M) at position 5 systematically abrogated TCR response.

For each of these combinations, we identified the sequences possessing the particular amino acid at the combined position and calculated the sequence-dependent diffRBM single-site factors for that position (Figure 3B). For all amino acid/position pairs, the distribution of these single-site factors is skewed toward positive values (Figures 3B, S6A), meaning that the diffRBM units predict a positive contribution to immunogenicity, and the degree of such contribution varies sequence by sequence in relation to the other amino acids. In contrast, measuring purely the amino acid frequency ratio at that position under the independent-site assumption predicts a contribution to immunogenicity that is significantly smaller (Figure 3B). In other words, the diffRBM’s ability to capture the sequence context reconciles its predictions with previous findings on residues that are key to immunogenicity. This result emphasizes that the understanding of the biochemical basis of immunogenicity can benefit from modeling approaches able to assess the role of single residues in a sequence context-dependent manner.

#### Model predictions are in agreement with data on TCR reactivity to mutant peptides

To further corroborate the connection between the model’s predictions and T-cell reactivity assays, we considered the data from [Łuksza et al., 2022] on the TCR response to one of the highly immunogenic peptides to which our HLA-A*02:01-specific diffRBM model of immunogenicity can be applied (NLVPMVATV from the human cytomegalovirus). These data measure TCR reactivity to all possible single-site mutants of NLVPMVATV for 3 TCRs specific to it (Methods, Figure 3C). Some of these single-site mutations do not cause a complete loss of TCR response (we refer to them as ‘non-lethal mutations’), meaning that TCR reactivity can be recovered by increasing the peptide concentration (Figure 3C). We can estimate a ‘cost’ for such mutations in terms of the TCR cross-reactivity between NLVPMVATV and its mutants, measured in [Łuksza et al., 2022] as the log ratio between the half maximal effective concentration for TCR activation after the peptide has been mutated 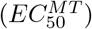 and the one before the mutation 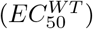, see Figure 3C. Positive costs reflect the increase in concentration needed to recover TCR reactivity and the corresponding decrease in peptide immunogenicity. Other mutations completely destroy peptide-TCR binding, and TCR reactivity cannot be restored even at highest peptide concentrations considered (we refer to them as ‘lethal mutations’, see Figure 3C).

Lethal mutations tend to occur at the typical TCR-contact positions along the peptide (Figure 3D), hence the diffRBM units, through the corresponding single-site factors, should account for the drop in TCR response upon these mutations. We found that the diffRBM units predict mainly positive costs for these mutations, which matches qualitatively the observed loss of immunogenicity (Figure 3E).

We observed that the magnitude of non-lethal mutations across all TCR-mutant pairs is concentrated at the peptide positions 2 and 9, which are the anchor sites for binding the HLA in HLA-A*02:01-specific peptides (Figure 3F). We therefore hypothesized that an increase in peptide concentration is needed first to compensate for decreased HLA presentability, hence a model of presentation alone should be capable of predicting their effect. We found that the prediction by our model of presentation by HLA-A*02:01 (background RBM) correlates significantly to the experimental mutation costs, and the degree of correlation is consistent across all 3 TCRs, given that these mutations disrupt predominantly peptide presentation on the HLA (Figures 3G, S6B). In contrast, the diffRBM units cannot predict these mutational effects concentrated at the anchor sites for presentation (Figure S6B), since its parameters capture the distinctive molecular composition of immunogenic peptides at the central positions (Figure 2C).

### 3.4 DiffRBM discriminates immunogenic *vs* non-immunogenic peptides

The diffRBM units learn distinctive sequence patterns of immunogenicity, having the background RBM captured the sequence constraints associated to presentability. Such patterns should contribute to distinguish immunogenic from non-immunogenic peptides. We therefore assigned scores of immunogenicity based on the diffRBM units to held-out test sets of peptides (Figure 4A) and we measured the score’s ability to discriminate HLA-specific immunogenic peptides from non-immunogenic ones sharing the same the HLA specificity in terms of the area under the receiver operating characteristic curve (AUC), see inset of Figure 4A and Methods.

**Figure 4:**
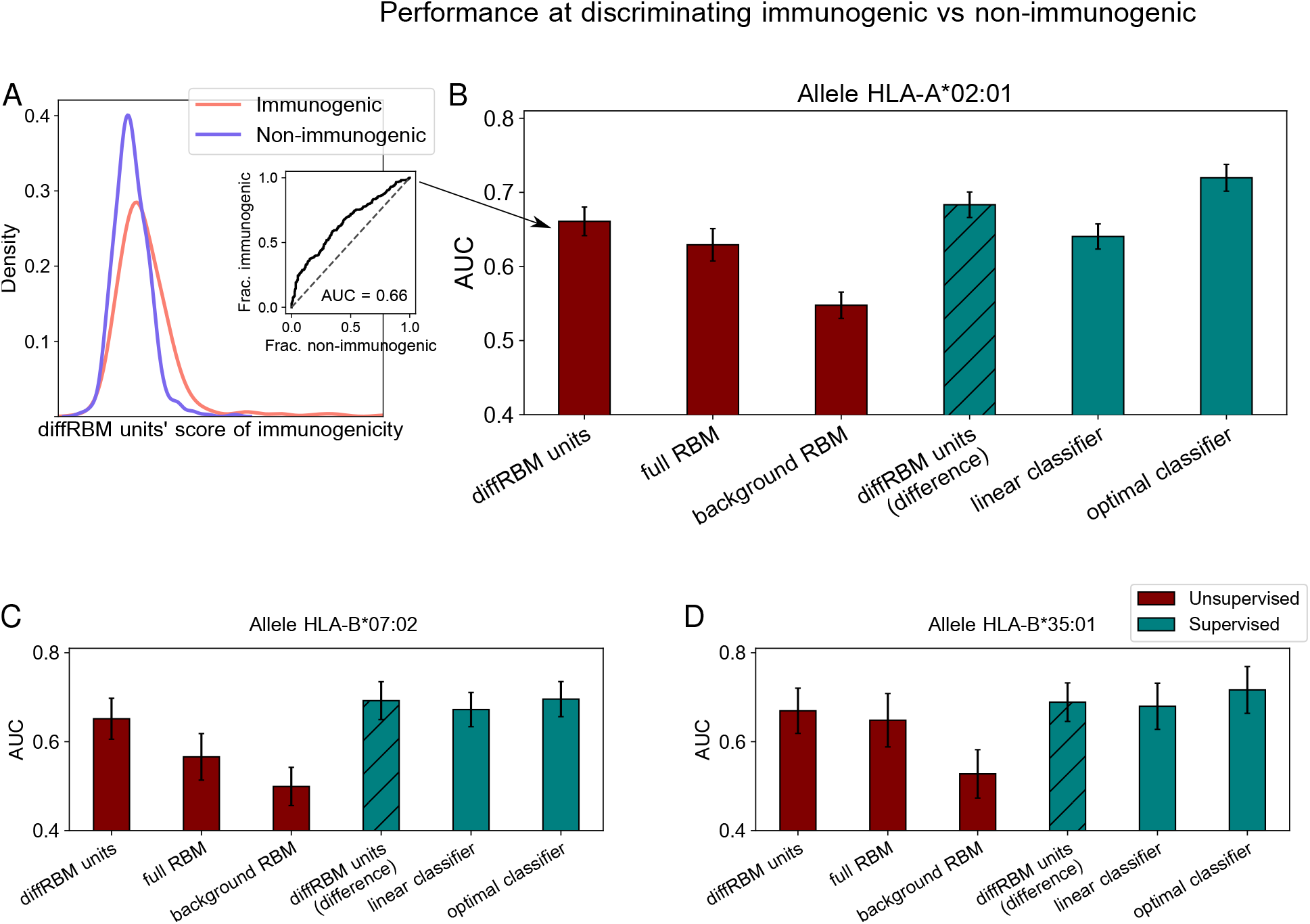
Performance at discriminating immunogenic *vs* non-immunogenic peptides. A: For a given HLA (*e.g*. HLA-A*02:01), we assign diffRBM units’ scores to held-out sets of immunogenic and non-immunogenic antigens and we measure the discrimination performance via the Area Under the Curve (AUC), see Methods. B: Performance of diffRBM units, full RBM, background RBM and other methods for the HLA-A*02:01-specific dataset as measured by the AUC. We have divided the methods into unsupervised and supervised ones (color code). Unsupervised methods are meant as the methods trained on immunogenic (or presented) only peptides to learn an unsupervised model. Supervised methods are trained on both immunogenic and non-immunogenic to optimally discriminate them. The approach ‘diffRBM units (difference)’ is intermediate: it exploits the annotation of peptides as immunogenic and non-immunogenic but it is not optimized for the discrimination task. C,D: Same plot as B for the immunogenicity models built for HLA-B*07:02 (C) and HLA-B*35:01 (D). AUC values shown are the average over 50 partitions into training and test sets and error bars give the corresponding standard deviation (Methods).

In Figure 4B, we compare the diffRBM units’ AUC for the HLA-A*02:01-specific model to the AUC of the full RBM, that includes the background model in its predictions and hence yields a joint representation of the enriched patterns and of the background constraints. The AUC of discrimination progressively decreases, as expected, from the model part that disentangles enriched patterns from the background constraints (diffRBM units), to the full RBM capturing both, to the model fit to the background constraints only (background RBM). (In fact, there is no reason why background RBM should predict anything at all unless presentability and immunogenicity are correlated). This trend indicates that learning the background constraints that are shared by immunogenic and non-immunogenic peptides along with the sequence pattern enrichment distinctive of immunogenicity can act as a confounding factor when we look for the features that characterize and distinguish immunogenic peptides. Also simpler differential models relying on the independent-site assumption, while returning lower AUCs than RBM-based models, exhibit a decrease in AUC between the differential part and the full model (Figure S7). The AUC values and trend remain stable when we compare scores assigned to immunogenic peptides and to peptides from the human proteome (Figure S8A) and when we score immunogenic and non-immunogenic peptides from the same organism having trained the models on the immunogenic peptides from all the other organisms (Figure S9).

Conversely, the diffRBM units are not designed to capture the background constraints (here associated to presentability). As a result, they cannot successfully discriminate presentable antigens from generic peptides that are predominantly non-presentable (like peptides randomly drawn from the human proteome), in contrast to background RBM and the full RBM (Figure S8B). The same trends of discrimination performance are consistently found across the 3 HLA alleles considered (Figures 4C-D, S7, S8A-B).

The prediction by the background model (trained on peptides probed for their binding to a given HLA-I) provides no clear signal that already the binding affinity to the HLA can discriminate immunogenic peptides (average AUC=0.55 for HLA-A*02:01, 0.50 for HLA-B*07:02, 0.53 for HLA-B*35:01, see Figure 4). To further check this prediction, we scored HLA-A*02:01 peptides by their binding affinity to the HLA through NetMHCpan4.1 [Reynisson et al., 2020] and found a comparable difference in score distributions between immunogenic and non-immunogenic peptides (AUC = 0.54 for HLA-A*02:01, 0.48 for HLA-B*07:02, 0.53 for HLA-B*35:01). Our observation is in line with a recent large-scale mapping of killer T-cell recognition of candidate neopeptides at high HLA affinity [Kristensen et al., 2022], which did not find a significantly different distribution of HLA-binding NetMHCpan scores between immunogenic *vs* non-immunogenic neopeptides. Other studies, however, have suggested that immunogenic peptides bind more strongly to the HLA compared to non-immunogenic ones, both in the case of viral epitopes [Croft et al., 2019, Buckley et al., 2022] and neo-epitopes [Bjerregaard et al., 2017, Buckley et al., 2022]. More work is needed in the future to clarify the link between binding affinity to the HLA and immunogenicity.

#### A deeper classifier reaches optimal performance, but diffRBM stays comparable

To perform a task of classification of immunogenic *vs* non-immunogenic peptides, it is more effective to leverage the information in both sets of immunogenic and non-immunogenic peptides. In our framework of differential learning, this can be done by training a second set of differential units on the non-immunogenic peptides, which learns differences in their amino acid statistics with respect to the background. The difference of the scores of the diffRBM units trained on immunogenic peptides and the ones trained on non-immunogenic peptides plays as well the role of a score of immunogenicity, expected to be positive when evaluated on immunogenic peptides and negative on non-immunogenic peptides (Methods). We use it to classify immunogenic *vs* non-immunogenic peptides, resorting to the AUC to measure its classification performance (Figure 4B-D). Such performance is, by construction, the same as the difference of the full RBM scores (Methods, Figure S10).

Exploiting the availability of both immunogenic and non-immunogenic peptides, we also trained a deep neural-network-based classifier. The classifier architecture was optimized among several ones of different depth and width (Methods, Figure S11, Table S2) and reaches a performance quantified by a cross-HLA average AUC = 0.71 ± 0.02 (see ‘optimal classifier’ in Figure 4B-D; here and in the following uncertainties are estimated over several training-test partitions, see Methods). As a term of reference we considered a linear classifier, whose performance (AUC = 0.66 ± 0.02) shows that there is a gain in resorting to a more flexible architecture in terms of depth and width. The AUC of the optimal classifier sets the maximal predictive performance that can be achieved, with the datasets under consideration, by a supervised method that is trained to discriminate immunogenic and non-immunogenic antigens. Its value (AUC = 0.71 ± 0.02) indicates that the predictability of immunogenicity from peptide sequences is limited, both by data availability and by the fact that sequence patterns along the peptide are not the only determinant of a positive T-cell response. In the future, more exhaustive assessments of peptide immunogenicity should account for the composition of cognate TCRs, peptide expression levels as well as the regulatory dynamics underlying T-cell response in physiological conditions. The performance of the diffRBM units’ scores (AUC = 0.69 ± 0.02) is slightly lower than the one of the optimal classifier but higher than the one of the linear classifier (Figure 4B-D). In this comparison, it is important to remark that the deep classifier is optimized for the classification task, while classification through the difference of diffRBM units’ scores is based on unsupervised models trained to learn the distribution of interest (the one of immunogenic or non-immunogenic antigens) and identify its distinguishing features from the baseline distribution of presented antigens.

Finally, we compared the discrimination performance of diffRBM to two established methods for immunogenicity prediction, the IEDB immunogenicity tool and PRIME [Schmidt et al., 2021]. Since these methods are not designed to be re-trained on custom datasets such as ours, the comparison cannot be fully consistent in terms of using the *same* training and test sets, as done for all the methods in Figure 4. The IEDB tool for immunogenicity prediction is based on the model by [Calis et al., 2013] and can be downloaded from http://tools.iedb.org/immunogenicity/. By applying it to our set of immunogenic and non-immunogenic peptides, we obtained AUC=0.54 for the HLA-A*02:01-specific peptides, AUC=0.60 for HLA-B*07:02 and AUC=0.57 for HLA-B*35:01, which are all lower than the diffRBM average AUC values (respectively 0.66, 0.65, 0.67, see Figure 4). By applying PRIME [Schmidt et al., 2021] to our set of immunogenic peptides, we found again poorer performance compared to the diffRBM units (average cross-HLA AUC = 0.51, see Methods), despite not having excluded from our test sets the immunogenic peptides that are either in PRIME’s or in the IEDB tool’s training sets. Differences in the peptide datasets used may contribute to explain this result. For instance, our dataset consists of epitopes with experimentally validated positive T-cell responses from IEDB, mainly of microbial origin, while PRIME’s training set [Schmidt et al., 2021] was constructed in such a way as to contain a high proportion of neoantigens.

As an additional point of comparison, Riley et al. [Riley et al., 2019] propose a neural network trained in a supervised way to classify immunogenic against non-immunogenic peptides using sequence as well as structural features of the peptide/HLA complex. When they check the performance of a peptide sequence-only predictor (using nonamer HLA-A*02:01-restricted peptides), they obtain AUC = 0.61 on the training set and AUC = 0.50 on the test. This further confirms that the performance of our diffRBM approach, despite not being directly trained as a discriminator of immunogenic *vs* non-immunogenic peptides, compares favorably to existing sequence-based tools of prediction.

### 3.5 DiffRBM model for T-cell specific binding

The idea of a differential learning of unsupervised models of TCR statistics, designed to describe the process of thymic selection acting on top of the receptor generation via V(D)J recombination, has first been put forward by [Elhanati et al., 2014]. This effort has culminated in the release of the software SONIA [Sethna et al., 2020] and, most recently, of its deeper version soNNia [Isacchini et al., 2021]. Depending on the TCR dataset used for training, these unsupervised models are applicable also as tools to classify TCRs by type (helper vs killer) and by epitope specificity [Isacchini et al., 2021]. The computational prediction of the epitope specificity of a given receptor, in particular, is extremely relevant for therapeutic design, and has been attempted through several algorithms [Cinelli et al., 2017, Gielis et al., 2019, Jokinen et al., 2019, Davidsen et al., 2019, Springer et al., 2020, Luu et al., 2021, Sidhom et al., 2021, Chronister et al., 2021, Weber et al., 2021, Lin et al., 2021, Montemurro et al., 2021, Zhang et al., 2021], spanning a variety of learning methods from random forests [Gielis et al., 2019] to deep neural network-like architectures (convolutional neural networks [Montemurro et al., 2021, Sidhom et al., 2021, Moris et al., 2021, Luu et al., 2021], bimodal attention networks [Weber et al., 2021], long short-term memory networks and autoencoders [Springer et al., 2020, Lu et al., 2021]). These models are typically calibrated towards achieving high predictive power as classifiers of TCR specificity, while less attention is paid to the interpretability of their predictions apart from a few exceptions [Papadopoulou et al., 2022]. Our diffRBM approach differs from and complements these existing methods because, similarly to SONIA, it focuses on the unsupervised discovery of salient amino acid patterns that can allow for biological insight into the molecular properties determining binding specificity.

To train diffRBM models of epitope specificity, we first collected from the VDJdb database [Shugay et al., 2018, Bagaev et al., 2020] datasets of TCRs specific to 4 epitopes (Methods): the M1_58_ peptide from the influenza virus (with sequence GILGFVFTL), the pp65_495_ from the human cytomegalovirus (CMV, with sequence NLVPMVATV), the BMLF1_280_ peptide from the Epstein-Barr virus (EBV, with sequence GLCTLVAML), the peptide from the Spike protein S_269_ from Sars-Cov-2 (with sequence YLQPRTFLL). We limited the search to sequences of the *β* chain of the TCR (TCR*β*), where the sites of binding to the antigen are concentrated in a region called complementarity determining region 3 (CDR3*β*).

The background dataset, in this case, is meant as a typical bulk TCR*β* repertoire in normal conditions (Figure S13). In particular we take, to train background RBM, the repertoire of a hypothetical universal donor that was constructed by [Isacchini et al., 2021] from the TCR*β* repertoires of the large scale study [Emerson et al., 2017] (Methods). The sequence features captured by background RBM concern germline-encoded amino acid usage related to stability and binding constraints (as is the case for the two conserved residues cysteine and phenylalanine delimiting the CDR3*β* region), as well as additional biases in amino acid usage stemming from VDJ recombination and thymic selection.

After having trained the background RBM, we train a set of diffRBM units on each set of epitope-specific CDR3*β* (Figures 5A, S13). By design, these diffRBM units capture antigen-driven convergent sequence features that have been documented in connection to epitope specificity [Dash et al., 2017, Glanville et al., 2017, Meysman et al., 2019, Pogorelyy and Shugay, 2019, Thakkar and Bailey-Kellogg, 2019, Mayer-Blackwell et al., 2021, Valkiers et al., 2021]. As such, similarly to the model of antigen immunogenicity, the diffRBM units can predict contact sites along the CDR3*β* (Figure 1E) and classify specific receptor against generic, predominantly non-specific ones (Figure 1F).

**Figure 5:**
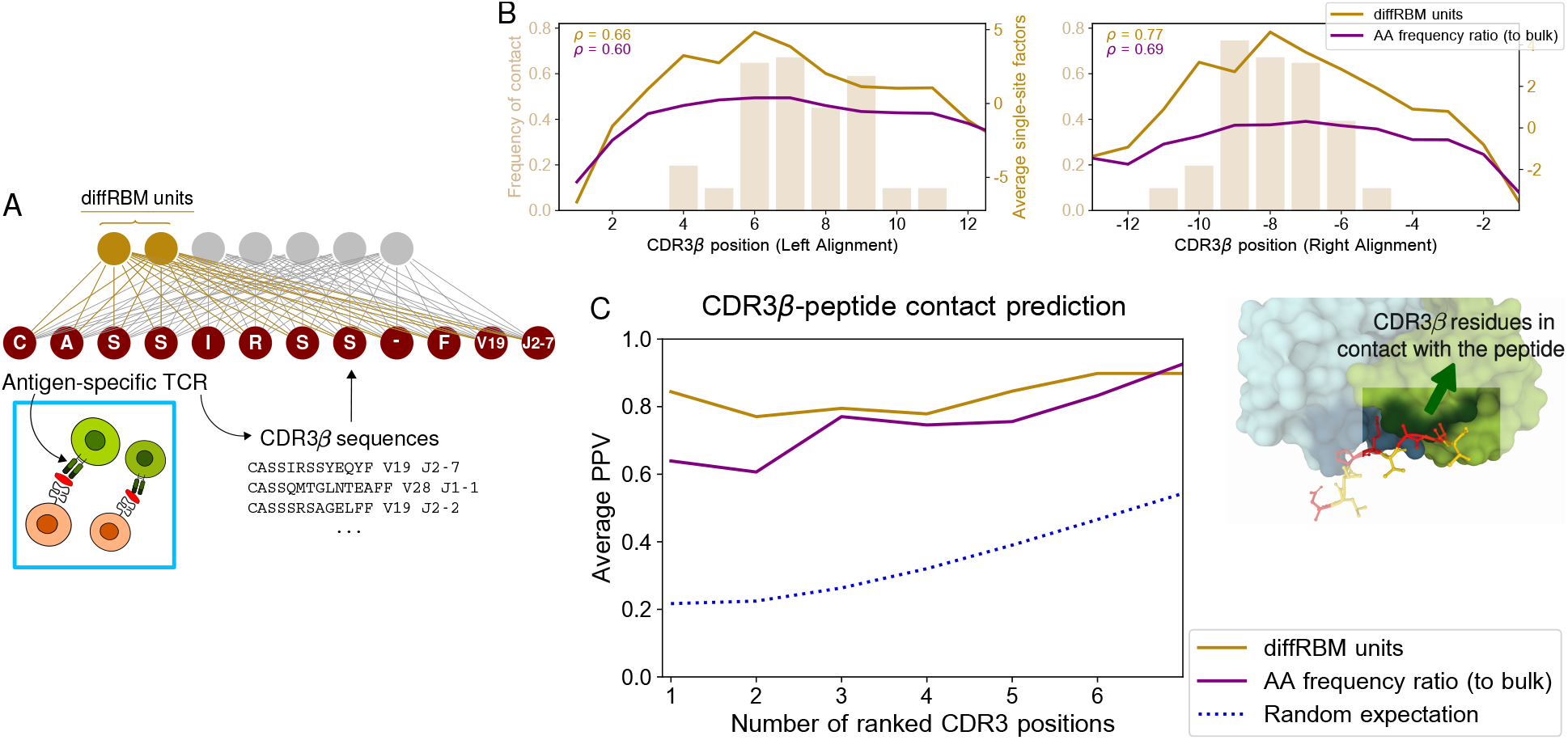
DiffRBM model of TCR epitope specificity and structural interpretation of its parameters. A: The diffRBM units are learnt from the CDR3*β* sequences of antigen-specific T-cell receptors. B: Frequency of contact with the peptide of each CDR3*β* position calculated across 12 structures (2 for YLQPRTFLL, 3 for NLVPMVATV, 1 for GLCTLVAML, 6 for GILGFVFTL). A position along the CDR3*β* is estimated as the distance either from the left or from the right anchor site. Contacts are estimated from the structures choosing a 4 Å distance cutoff between CDR3*β* and peptide. We plot, along with the contact frequency distribution, the magnitude of single-site factors based on the diffRBM units or the amino acid frequency ratio (of peptide-specific sequences relative to bulk-repertoire sequences) averaged over the 12 CDR3*β*. C: Average PPV for the prediction of CDR3*β* positions in contact with the peptide as a function of the number of ranked positions (the average is calculated over the 12 structures listed in the caption of B). For each CDR3*β* sequence, predictions are obtained from the single-site factors of the corresponding peptide-specific models of TCR binding (diffRBM or the model based on the amino acid frequency ratio). The PPV is reweighed by a term accounting for the CDR3*β* sequence similarity (Methods, Figure S15A-B).

### 3.6 DiffRBM predicts CDR3*β* contact positions with the peptide

It has been discussed that convergent features in receptors responding to same antigens have structural interpretation in terms of interactions across the peptide-TCR interface [Dash et al., 2017, Glanville et al., 2017] and that the TCR contact regions are dominated by CDR3*β* residues [Glanville et al., 2017, Ostmeyer et al., 2019, Milighetti et al., 2021] (see also Figure 1E). Starting from the available TCR-pMHC structures already analyzed, we looked at the peptide-TCR contacts lying in the CDR3*β* region of the TCR, focusing on the structures involving the 4 epitopes under consideration (12 structures in total, see Methods). Figure 5B shows the distribution of contacts along the CDR3*β* positions, which consists, as already observed in [Glanville et al., 2017, Ostmeyer et al., 2019, Milighetti et al., 2021], of stretches of 3-5 contiguous amino acids in the central part of the CDR3 (6-8 positions from the left and right anchor).

We estimated the diffRBM single-site factors for the CDR3*β* sequences from the available structures. Their average value across CDR3*β* sequences concentrates on the CDR3*β* central positions and well correlates with the contact frequency distribution (Figure 5B). We next ranked the positions by the singlesite factor magnitude and we took top ranking positions as positions of predicted contact. Figure 5C shows the PPV averaged over the 12 available structures (Methods). Similarly to the predictions for peptides (Figure 2D), the predictive power of the diffRBM units is superior to the position-specific amino acid frequency ratio between the antigen-specific and the bulk-repertoire set of receptors. The PPV trend stays robust varying the distance cutoff from 4 up to 5 Å (the value chosen to determine peptide-TCR contacts in other work [Calis et al., 2012, Glanville et al., 2017, Ostmeyer et al., 2019, Milighetti et al., 2021]), see Figure S15D.

### 3.7 DiffRBM discriminates specific receptors

We tested the power to discriminate receptors specific to the 4 epitopes under consideration (GILGFVFTL, NLVPMVATV, GLCTLVAML, YLQPRTFLL) from generic sequences drawn from a bulk repertoire (background dataset). These can be seen as a proxy for non-specific receptors, since the large majority of them is not expected to respond to a specific epitope. We measured the performance at discriminating specific from generic receptors by the AUC (Figure 6A) and the results for the 4 epitope-specific models are reported in Figures 6B-E, S16. We observed the same trend described for the models of immunogenicity, whereby the AUC of discrimination for the diffRBM units is consistently higher than the one for the full RBM. Also in this context, singling out the sequence features associated to epitope specificity, as the differential units do, enhances the model’s predictive performance compared to the full RBM, where the information on those features is added to the background constraints. Any discrimination power is lost when using the background RBM, as it should, since it has no information on epitope specificity.

**Figure 6:**
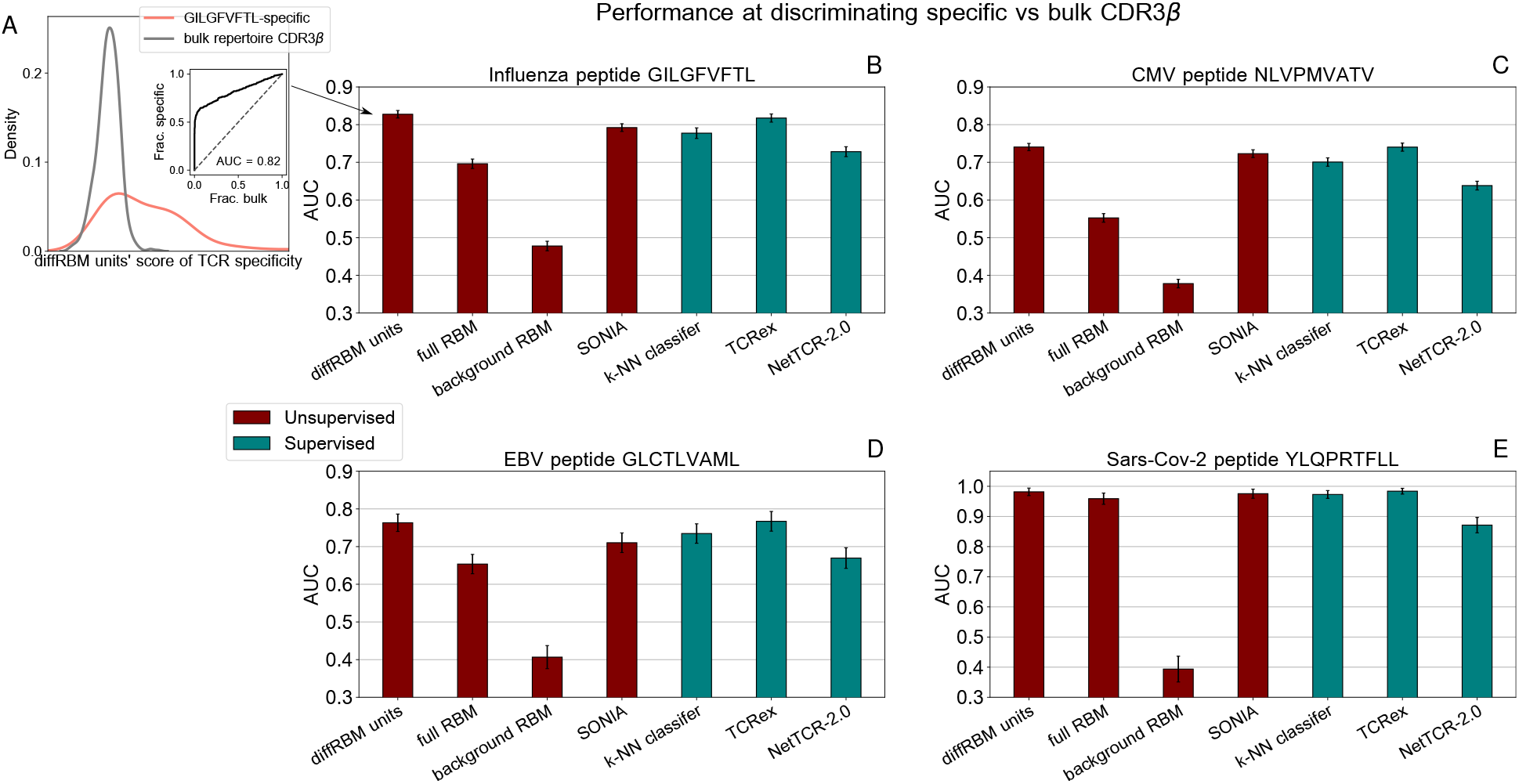
Performance at discriminating antigen-specific from generic T-cell receptors. A: For a given epitope model (*e.g*. the Influenza epitope GILGFVFTL), we assign diffRBM units’ scores to held-out sets of antigen-specific CDR3*β* and generic CDR3*β* from the bulk repertoire, and we measure the discrimination performance via the Area Under the Curve (AUC), see Methods. B: AUC of the diffRBM units, full RBM, background RBM and other supervised and unsupervised methods trained and tested on CDR3*β* sequences specific to the Influenza epitope GILGFVFTL. C-E: The performance assessment illustrated in A-B is repeated for the models of specific response to the CMV epitope NLVPMVATV (C) EBV epitope GLCTLVAML (D), and the Sars-Cov-2 epitope YLQPRTFLL (E). AUC values shown are the average over 50 partitions into training and test sets and error bars give the corresponding standard deviation (Methods).

To further check the robustness of our model’s performance, we also repeated the training and testing constructing the background model from a different set of bulk-repertoire TCR*β* sequences from healthy donors (the dataset from [Dean et al., 2015]). All the performance metrics are almost unaffected by this change of the background dataset (Figure S17). The average AUC attained by the diffRBM units across the 4 peptide-specific models is 0.83 ±0.01 with the background dataset from [Emerson et al., 2017] and 0.82±0.01 with the background dataset from [Dean et al., 2015] (uncertainties are estimated over several training-test partitions, see Methods).

#### DiffRBM reaches state-of-the-art performance

For the sake of comparison of diffRBM to other tools, we considered another generative model of antigen-specific repertoires, SONIA [Elhanati et al., 2014, Sethna et al., 2020], and a series of methods that are trained to discriminate target-specific from unspecific receptors (labeled as ‘supervised’ in Figure 6B-E). First, as baseline supervised method to compare with, we considered a *k*-Nearest Neighbours (*k*-NN) classifier (Methods). In [Weber et al., 2021] a *k*-NN classifier served as a baseline method to determine the predictive power achievable from TCR sequence similarity alone and, despite its simplicity, it was found to outperform some existing methods. Next, we compared diffRBM to state-of-the-art predictors of TCR specificity, namely TCRex [Gielis et al., 2019] and NetTCR-2.0 [Montemurro et al., 2021]. TCRex is a random forest based classifier of TCRs by epitope specificity, while NetTCR belongs to a family of methods [Springer et al., 2020, Weber et al., 2021, Milighetti et al., 2021] that are trained on TCR-antigen pairs to predict a score of binding, but can also be applied to the prediction of whether an unseen TCR is likely to be specific to a given peptide target.

To make a fair comparison of performance, we re-trained all these methods on our training sets and evaluated the AUC of discrimination of specific receptors on our test sets. For TCRex and NetTCR-2.0, we considered their version for the *β*-chain only (Methods). The performance of the diffRBM units is fully comparable to TCRex (both with an average AUC across peptides of 0.83 ± 0.01) and only slightly higher than SONIA and *k*-NN (AUC = 0.80 ± 0.01 for both), see Figure 6B-E. We observed a moderately lower performance of NetTCR-2.0 (AUC= 0.73 ± 0.01), which could be partially due to the fact that the CDR3*β* version of NetTCR-2.0 does not take as input the V and J genes. To check the role of including information on V and J, we also re-trained diffRBM using only the CDR3*β* amino acid sequence (Figure S19). The performance of the diffRBM units is reduced (AUC = 0.79 ± 0.01), but still higher than NetTCR-2.0. We note, however, that the strength of methods like NetTCR [Montemurro et al., 2021, Springer et al., 2020, Weber et al., 2021, Milighetti et al., 2021] relies on being trained to be ‘pan-specific’, that is, predictive across potentially any peptide and not only for specific peptides. An approach like diffRBM (similarly to SONIA) returns peptide-specific models and hence needs to be trained on TCRs known to recognize a certain peptide for any prediction of specificity to that peptide. Furthermore, the performance by diffRBM decreases compared to other methods when we look at a different task, the one of discriminating a set of antigen-specific receptors from receptors with another antigen specificity [Meysman et al., 2022]. This is due to the fact that our approach, in contrast to other methods [Gielis et al., 2019, Montemurro et al., 2021, Weber et al., 2021], is unsupervised, meaning that it is not trained using receptors of different antigen specificity as negatives against which the positives should be discriminated.

Overall, these comparisons support the conclusion that our approach, when used for classifying receptors that are epitope binders *vs* generic non-binders from a bulk repertoire, reaches the performance of state-of-the-art TCR specificity predictors [Gielis et al., 2019, Montemurro et al., 2021, Isacchini et al., 2021].

## 4 Discussion

We have introduced a framework, based on the probabilistic graphical model of Restricted Boltzmann Machines, aimed at learning what we called ‘differential’ models, which capture differences of selected subsets of sequences with respect to the full set of sequences of similar type (the ‘background’). More specifically, the ‘differential’ part of the model (the diffRBM units) is designed to learn such differences (Figures 1, S1). First, we applied this framework to the problem of modeling antigen immunogenicity, the ability to trigger a positive T-cell response. We showed that the diffRBM units are able to encode several features relevant to immunogenicity. More precisely, we used the parameters associated to the diffRBM units to estimate which positions are more likely to be in contact with the TCR and we showed that they can distinguish immunogenic from non-immunogenic peptides better than a full RBM, their performance being comparable to a supervised model (the ‘optimal classifier’) that we built and optimized for classification.

To work with samples of sufficient size, the diffRBM models of immunogenicity discussed here were trained on IEDB peptide data, which are predominantly pathogenic. The diffRBM approach is in principle applicable also to cancer neoantigen discovery, but the particular models may need to be retrained on neoantigen-specific training sets, given that substantial variation in method performance and in the immunogenicity-related predicted features between pathogenic and cancer data has been evidenced [Buckley et al., 2022]. We did an additional analysis as a preliminary check on the diffRBM performance in the cancer setting: we assigned scores to the HLA-A*02:01-presented neoantigens from the TESLA dataset [Wells et al., 2020] (11 immunogenic and 227 non-immunogenic peptides) and we obtained an AUC of immunogenic *vs* non-immunogenic discrimination differing significantly from the random expectation (0.6 for the diffRBM units’ scores and 0.69 for the difference of diffRBM units’ scores).

We also learnt differential models from datasets of epitope-specific T-cell receptors to capture the sequence features underlying their ability to bind to a given epitope, and we have successfully tested the models’ power to discriminate epitope-specific from generic receptors and to identify CDR3*β* residues in contact with the antigen. We looked at the task of distinguishing a small number of antigen-specific receptors from the bulk repertoire (forming the background), a task which reflects many typical scenarios of application. Indeed, antigen-specific receptors constitute a minority of the repertoire, with experimental estimates of the fraction of the TCR repertoire reactive to a given epitope ranging between 10^−6^ and 10^−4^ [Yates, 2014]. It is at this task that diffRBM performs as well as the state-of-the-art methods. The performance by diffRBM decreases when we consider as negatives receptors with a different antigen specificity [Meysman et al., 2022], because the differential units can learn features shared between antigen-specific sequences (*e.g*., being expressed by CD8+ T cells). More fine-tuned choices of the background (*e.g*., with only CD8+ T-cell receptors) would be needed to improve performance for this particular task.

The diffRBM model of specific TCR response can be formulated in an equivalent way for the CDR3 region on the other chain of TCRs (*α* chain). In this case, one uses CDR3*α* sequences from healthy TCR repertoires and samples of antigen-specific CDR3*α* as respectively the background and the selected dataset. DiffRBM models for the *α* chain reach a discrimination performance comparable to the one for the *β* chain, as shown in [Meysman et al., 2022]. On the same footing, with single-cell TCR sequencing data becoming increasingly available, our approach could also be extended to modeling the pairs of TCR *α* and *β* chains, which have been suggested to play a synergistic role in determining antigen specificity [Carter et al., 2019, Montemurro et al., 2021, Milighetti et al., 2021].

Our primary goal was to learn, through diffRBM, generative models that can describe the probability distribution of data with ‘selected’ features compared to ‘background’ data. As such, they can be used to calculate a probabilistic measure of the diversity of the space sampled by the data, the entropy of the inferred probability distribution (Methods), to then estimate the change in diversity due to selection. The entropy values of background and selected sequences obtained through our models of immunogenicity and TCR specificity highlight a reduction of diversity in selected sequences stemming from the enrichment in distinctive sequence patterns (Figure S20). Such a reduction has substantial variability across epitopes in the case of TCR specificity models (Figure S20B), reflecting the existence of more or less well-defined sequence motifs underlying epitope specificity.

In this paper, the application of our approach has been clearly limited by the scarcity of experimentally confirmed immunogenic peptides for the majority of HLA alleles, as well as TCR specific to many other epitopes. Even for the HLA alleles and epitopes we considered, available datasets cover only a small fraction of HLA-specific peptides that have the potential to be immunogenic and TCR that can recognize given peptides, and pool together data obtained through a heterogeneous set of T-cell response assays, a source of additional noise in the data (Methods). Such limitations in the datasets might affect the performance of all the methods we tested. However, we emphasize here the methodological novelty of learning differential models of immunogenicity and antigen specificity. Our study provides robust proof-of-concept results that a differential modeling approach is an effective tool to extract the sequence features underlying the specificity of binding between antigens and T-cell receptors, and to generate insight into the structural basis of this process by identifying putative contact sites from the sequence alone. Furthermore, relying on unsupervised learning has the advantage that negative samples, whose construction is somewhat arbitrary, are not needed. In addition our model is generative, *i.e*., it could a priori be used to produce new, putative sequences of antigens or TCR with desired properties, testable in experimental setups.

The impact of biochemical properties, such as hydrophobicity, on immunogenicity is mediated by the structural arrangement of the peptide within the HLA binding groove and of TCR chains, hence structural features associated to immunogenicity have been shown to be important for predictive power, both at the level of peptide binding to the MHC [Riley et al., 2019] and TCR binding to the pMHC complex [Lin et al., 2021]. Recent work [Lin et al., 2021] offers an approach for predicting immunogenic peptides by incorporating structural information into the estimation of energies of TCR-antigen binding using coarse-grained protein folding models. Its use is essentially limited to TCR-peptide pairs of known crystal structures, narrowing down the scope of its application. The work [Riley et al., 2019] proposes a neural network-based predictor of immunogenicity that uses the peptide sequence as well as information on the peptide/MHC complex (such as van der Waals interactions, Coulomb potentials, hydrogen bond energies, solvent accessible surface areas) estimated by rapid structural modeling with Rosetta [Chaudhury et al., 2010]. There the need for peptide/MHC structure is circumvented by threading generic peptide sequences into pre-defined template structures, a strategy which is successful due to the homogeneity of the backbone conformation across nonamer peptides presented by the same HLA-I, but might become problematic with peptides of different length or for HLA types admitting different binding modes with the peptide [Gfeller et al., 2018]. Given the large availability of peptide sequence data, many of them produced in clinical settings for personalized medicine purposes, it is extremely valuable to develop approaches that can achieve the more broadly applicable and computationally faster task of scoring peptides by their potential to be immunogenic given the peptide sequence alone. Nevertheless, in terms of future work, leveraging structural information on the TCR-pMHC complex and its estimated binding energy along with sequences, as investigated by [Riley et al., 2019, Lin et al., 2021, Milighetti et al., 2021, Karnaukhov et al., 2022], will contribute to improving the modeling strategy presented, especially given the expected increase in the number of available crystallographic structures. These tasks will require *ad hoc* adjustment of the model architecture, to integrate structural and energetic features into its input, and hence goes beyond the scope of the more general and flexible scheme we aimed to proposed here, which relies on a ‘background’ dataset of large size, and a small subset of selected data with distinctive property that justify their selection. Such flexibility suggests its broad applicability in several contexts, such as SELEX experiments [Ellington and Szostak, 1990, Tuerk and Gold, 1990, Sola et al., 2020], where each round performs a selection of a subset of molecules from the previous, more populated round (the ‘background’).

## 5 Materials and Methods

### 5.1 DiffRBM architecture

The core idea of what we refer to as a ‘differential’ probabilistic model is to learn a distribution for sequence data ***σ*** with the parametric form:

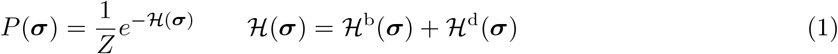

where *Z* is simply a normalization factor, 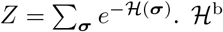 specifies the background distribution *P*^b^ through 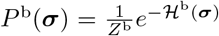, where 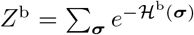. The background distribution *P*^b^(***σ***) is inferred from what we call a ‘background’ dataset 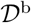, a large-size dataset that enables *P*^b^ to embed the main statistical properties of the data of interest with high accuracy (thanks to its large size). On the other hand, 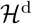 contains the parameters that are learnt on top of the background distribution from a set of ‘selected’ data 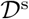, picking up their statistical differences with respect to the background distribution.

While this formulation allows for a rather generic choice of both 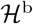 and 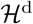, here we assume that they are parametrized in terms of Restricted Boltzmann Machines [Hinton, 2002, Hinton and Salakhutdinov, 2006]. Hence *P*^b^(***σ***) can be written as:

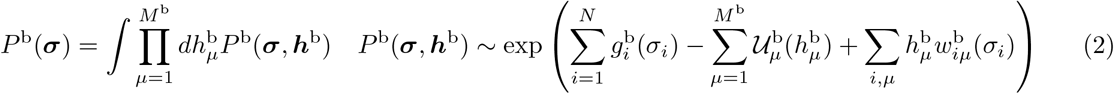

In a Restricted Boltzmann Machine, the probability of the data *P*^b^(***σ***) is expressed as the marginal of a joint probability over the data ***σ*** (the ‘observed’ sequences of length *N*) and a set of *M*^b^ ‘hidden’ units ***h***^b^, playing the role of coordinates of low-dimensional representations of the data. The joint probability is specified by a set of parameters whose value is inferred from the data (here protein sequences), consisting of: a set of single-site biases 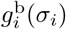, capturing the amino acid usage at each sequence position, a potential 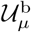 for each hidden unit 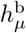 and a set of parameters 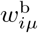, called weights, connecting the sites of observed sequences to each hidden unit. Thus for an RBM as background one obtains:

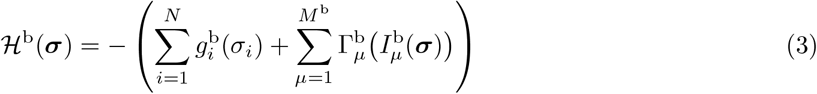

where we have set 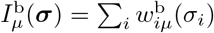 and 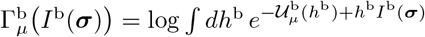. The parameters 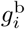 and 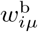, as well as the parameters defining the potentials 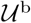, are learnt from the background dataset 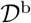 by maximizing the log-likelihood:

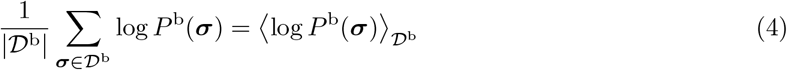

After the background distribution has been learnt, we learn the ‘differential’ units of the architecture that we have named differential Restricted Boltzmann Machine (diffRBM), corresponding to the probability distribution:

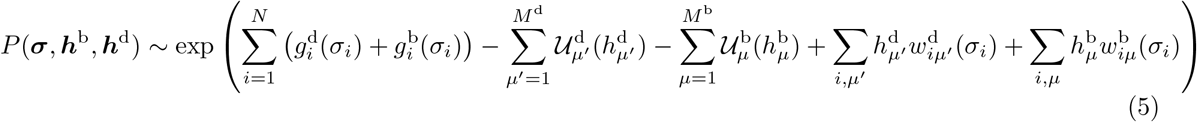

see also Figure S1. Hence:

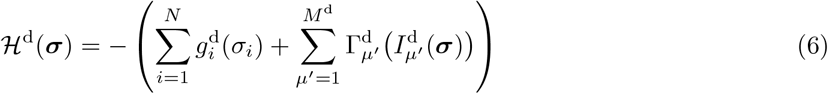

where we have defined 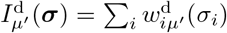 and 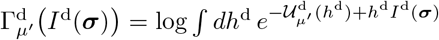. The parameters featuring in 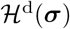 and defining the ‘differential’ RBM units are learnt from the dataset 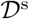 by maximizing:

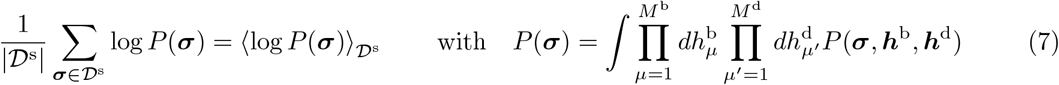

where *P*(***σ***, ***h***^b^, ***h***^d^) is given by (5). Let us define a compact notation **Ψ**^d^ for the set of parameters associated to the differential units, 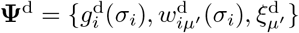, where 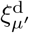 is a shorthand for the parameters specifying the shape of 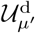 (in all our implementations, both 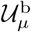 and 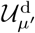 are set to dReLU potentials, a choice that has been shown to confer high expressivity [Tubiana et al., 2019]). The rules to infer **Ψ**^d^ are given by gradient-ascent equations to maximize the likelihood of the post-selection data 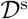 under the full RBM model (equation 1):

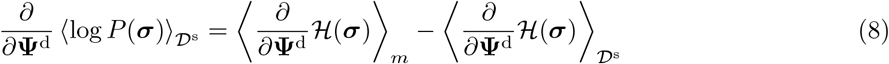

where 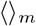 indicates the average under the full RBM model, 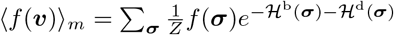. Since the parameters **Ψ**^d^ to learn appear only in 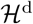, one has 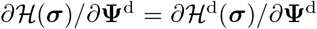; the contribution of the background 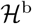 enters therefore only in the estimation of the average 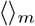, which requires, at each step of the training, to sample configurations with the probability (1).

In summary, the diffRBM architecture (equation 5) is essentially equivalent to an RBM with *M*^b^+*M*^d^ hidden units and where the overall observed fields are written as 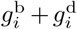 the weights 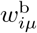 and potentials 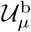 for the first *M*^b^ units (as well as 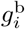) are learnt from the background data, then they are kept fixed and the weights 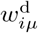 and potentials 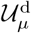 for additional *M*^d^ units (as well as 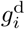) are learnt from the selected data (Figure S1).

Regularization terms can be added during the training to control the values of the inferred parameters. Here we used a *L*_1_-type regularization over the weights, which enforces sparsity to prevent overfitting (see [Tubiana et al., 2019] for details); following the convention in [Tubiana et al., 2019] we denote by 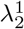 the coefficient setting the magnitude of such regularization term. In addition, a *L*_2_-type regularization on the fields was used for all models to control for large values stemming from undersampling, and following a standard choice [Cocco et al., 2018] its magnitude was set to the inverse of the size of the training dataset (hence to 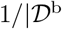 for background models and 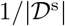 for differential models).

The predictions of peptide immunogenicity or epitope specificity rely on the assignment to sequences ***σ*** of scores. Using (1), the score of the full RBM is given by the log-likelihood:

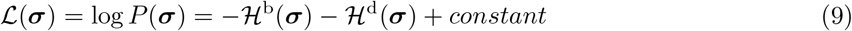

where 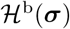 and 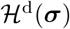 are given by (3) and (6) and the *constant* stands for a sequence-independent term coming from the partition function. Analogously, the background RBM score and the diffRBM units’ score are respectively:

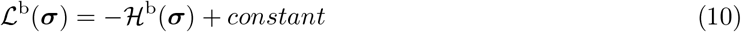

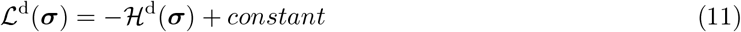

The entropies of background RBM and full RBM were estimated respectively as:

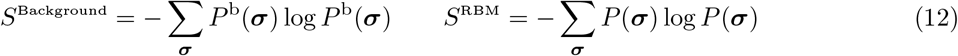

where log is meant as a natural logarithm, hence the entropy values in Figure S20 are expressed in nats.

In terms of software implementation, diffRBM is coded via a series of additional functions to execute a differential learning on top of the RBM Python implementation from [van der Plas et al., 2022] and is available at github.com/cossio/diffRBM. The codes used for its application to modeling antigen immunogenicity and TCR specificity are downloadable from github.com/bravib/diffRBM_immunogenicity_TCRspecificity.

### 5.2 Data collection

#### 5.2.1 Sequence datasets for the immunogenicity model

The differential models to describe immunogenicity were trained on sets of immunogenic peptides collected from the Immune Epitope Database (IEDB) [Vita et al., 2019], where the database entries were filtered through the following steps. Firstly, the curated set of HLA ligands tested in T cell assays was downloaded from IEDB (file *tcell_full_v3.csv* from http://www.iedb.org/database_export_v3.php, accessed in December 2021). We selected from this file linear, human peptides with a given HLA restriction (*e.g*. HLA-A*02:01), limiting the search to peptides of length 8-11 amino acids like in [Bravi et al., 2021b] and presented by HLA of class I (*i.e*., targeted epitopes of killer T cells). Following [Calis et al., 2013], we required the peptide (and not the full protein or the pathogen) to be the first immunogen (by setting the field *Antigen Epitope Relation* = ‘Epitope’) and we excluded T-cell response experiments with a restimulation step (by discarding ‘Restimulation in vitro’ from the field *In Vitro Process Type*). Immunogenic peptides were finally identified as the peptides for which positive responses by T cells were reported while negative ones were absent (field *Qualitative Measure* marked as ‘Positive’ or ‘Positive-High’ and never as ‘Negative’). Equivalently, non-immunogenic peptides were identified as the peptides for which negative responses by T cells were reported while positive ones were absent (field *Qualitative Measure* set to ‘Negative’ and never to ‘Positive’, ‘Positive-High’, ‘Positive-Intermediate’ or ‘Positive-Low’). To avoid oversampling, we removed duplicate entries. To check whether we needed an additional redundancy filtering, similarly to [Calis et al., 2013], we applied a reweighting scheme [Morcos et al., 2011] that reweighs each sequence by the inverse of the number of other sequences that have more than 80% of similarity, and we found that the models’ performance (Figure S12) is largely unchanged compared to the one without reweighting (Figure 4), indicating that there is no substantial need for additional sampling bias mitigation strategies. We choose only the HLA-I alleles for which the filtering steps just described allowed us to recover at least 200 immunogenic peptides and for which at least one TCR-pMHC structure was available in the Protein Data Bank (resulting in the choice of HLA-A*02:01, HLA-B*07:02 and HLA-B*35:01). The size of the final datasets of immunogenic peptides is: 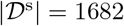 for HLA-A*02:01, 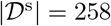 for HLA-B*07:02, 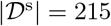 for HLA-B*35:01. Sets of non-immunogenic peptides consist of 2301 sequences (HLA-A*02:01), 807 (HLA-B*07:02), 166 (HLA-B*35:01).

To train the antigen presentation model, playing here the role of the background model, we relied on the sets of 8-11 amino acid long peptides extracted from IEDB by the RBM-MHC algorithm as described in [Bravi et al., 2021b], choosing the option of peptides from HLA binding affinity assays rather than mass spectrometry, to avoid biases in the amino acid statistics of presented antigens that might be due to mass spectrometry techniques. The resulting training dataset sizes for the background models are 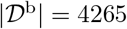 for HLA-A*02:01, 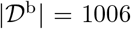 for HLA-B*07:02, 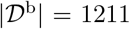 for HLA-B*35:01. For consistency with the type of datasets used to train the presentation model by RBM-MHC, scores of presentation from the algorithm NetMHCpan4.1 [Reynisson et al., 2020] are obtained with the option *-BA* (predictions from the training on binding assay data).

#### 5.2.2 Sequence datasets for the T-cell specificity model

Each differential model of specific T-cell response to a given peptide was trained on sets of T-cell receptors experimentally validated to be specific to the peptide, which were collected from the VDJdb database [Shugay et al., 2018, Bagaev et al., 2020] (file *vdjdb.txt* downloaded from https://vdjdb.cdr3.net in July 2021). More precisely, in the database we selected all the human TCR*β* chains fully annotated with their V and J segment and labelled to be specific to the given peptide (for example, for the Influenza M1_58_-specific model we set *antigen.epitope* = ‘GILGFVFTL’). We constructed the final training sets from the CDR3*β* sequence, the V and J segments identity of these entries, removing replicates with the same CDR3*β* and V/J annotation. We obtained datasets of epitope-specific receptors of size 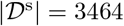 (for the Influenza M1_58_ model), 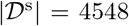 (for the CMV pp65_495_ model), 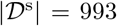 (for the EBV BMLF1_280_ model), and 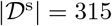 (for the Sars-Cov-2 S_269_ model).

For the background model (here meant as a model of the normal bulk TCR*β* repertoire), we considered: the dataset assembled by [Isacchini et al., 2021] pooling together unique TCR*β* clones from the 743 donors of the cohort in [Emerson et al., 2017], with a total of ~ 9 × 10^7^ sequences; the dataset of TCR*β* sequences from healthy donors from [Dean et al., 2015], already preprocessed by [Isacchini et al., 2021] and composed of ~ 2 × 10^6^ sequences. For training the background RBM on these data, we used smaller datasets of 10^6^ sequences that could be more easily handled by randomly subsampling from the entire datasets from [Emerson et al., 2017] and [Dean et al., 2015].

#### 5.2.3 Data preprocessing and formatting

RBM and PWM-based approaches require sequence inputs of fixed length, hence we performed an alignment. Background datasets are aligned to obtain same-length sequences, following the alignment procedures described in [Bravi et al., 2021b] for peptides and [Bravi et al., 2021a] for CDR3*β* amino acid sequences. The length of the alignment is set to 9 (9 being the typical length of HLA-I ligands) and to 20 in the case of CDR3*β*. These alignments serve as seeds to learn Hidden Markov Model profiles of length 9 and 20, in such a way that the selected datasets can be aligned against the profile built from the corresponding background dataset (see [Bravi et al., 2021b] for more details). In the models for TCR sequences, as a default, the input combines the aligned CDR3*β* amino acid sequence to the V segment type and the J segment type, all converted into numerical values varying within an interval of appropriate length (length = 21 for the CDR3*β* positions, standing for the 20 amino acids + 1 gap; length = 48 for the V type, length = 13 for the J type).

#### 5.2.4 Crystallographic structures from PDB

We downloaded, from the Protein Data Bank www.rcsb.org [Berman et al., 2000] as of February 2022, the TCR-pMHC crystallographic structures with 9 amino acid-long peptides where the HLA complex is HLA-B*35:01, HLA-A*02:01, or HLA-B*07:02. We excluded the structures with modified/non-peptidic epitopes and with incomplete TCR chains. As a result, we obtained 5 structures for HLA-B*35:01, 56 for HLA-A*02:01, and 1 for HLA-B*07:02. Some of the 56 HLA-A*02:01 structures describe TCRs in contact with the peptides we considered for the differential models of specific TCR response (3 for the Sars-Cov-2 epitope YLQPRTFLL, 3 for the CMV epitope NLVPMVATV, 1 for the EBV epitope GLCTLVAML, and 8 for the Influenza epitope GILGFVFTL).

For each structure, we estimated the positions along the peptide in contact with the TCR, using a standard cutoff at 4 Å [Rossjohn et al., 2015, Schmidt et al., 2021, Lu et al., 2021] between heavy atoms. The availability of structures is highly skewed toward the limited set of epitopes that have been the focus of several studies, hence our final list of peptides has some redundancy, with same or similar peptides in complex with different TCRs. If more than 1 structure contain the same peptide and have same contact positions, we retain only one of such structures (resulting in 4 structures for HLA-B*35:01, 41 for HLA-A*02:01, and 1 for HLA-B*07:02). On the other hand, if the same ligand is annotated with different contact positions, we keep these as different entries but we re-weight their contribution to the average PPV and the frequency of contact positions (section 5.7). We followed the same steps to estimate CDR3*β* contacts with the peptide and peptide contacts with the HLA complex and to filter out redundant entries, opting for a slightly more restrictive cutoff distance (3.5 Å) in the case of peptide-HLA contacts. Since distance cutoffs can vary with the van der Waals’ radii for single atoms [Sheriff et al., 1987], we also monitored the robustness of our results to changes in the choice of the cutoff (Figures S4D, S5D, S15D). The list of all structures and corresponding estimated contacts is provided in Table S1.

### 5.3 DiffRBM training and model selection

The first step of the diffRBM training consists of training the background model on the background dataset. For the model of immunogenicity, we trained allele-specific presentation models with an RBM architecture by running the RBM-MHC algorithm [Bravi et al., 2021b] with default parameters (10 hidden units, 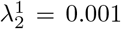). The peptide data used by default as training set by RBM-MHC to build allele-specific presentation models are downloaded from IEDB, see [Bravi et al., 2021b]. The RBM-MHC algorithm internally aligns peptide sequences of its training dataset to the reference length of 9 amino acids; we utilized the same alignment routine to align the immunogenic peptides against the seed given by the RBM-MHC training data.

The second step is training the diffRBM units on the ‘selected’ datasets, here immunogenic peptides and antigen-specific CDR3*β*s (section 5.2). Note that, for the applications discussed in this paper, the training on the background dataset and the one of the selected dataset are implemented *consecutively*. While the diffRBM architecture allows for a parallel training, the appeal to the sequential training here is justified by the size of the datasets at out disposal 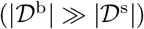 and by the particular problem under consideration (modeling an enrichment of specific features, occurring on top of baseline features, which do not change as feedback from the former). The datasets 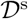, collected as described in section 5.2, are divided into a training set with 80% of the data (used for training and for model selection) and a test set with the remaining 20% (used for model validation, see section 5.5), repeating this split 50 times.

Having fixed the background model, we used the largest available dataset of immunogenic sequences (the one for HLA-A*02:01) to perform model selection by cross-validation, as follows. We further divided randomly each of the 50 training sets into a set actually used for training and a validation set (with respectively 80% and 20% of the training set). We used this training/validation partitions to select optimal hyperparameters for the differential part (number of the number of hidden units and regularization penalty 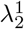), by training diffRBM models on the training set at varying hyperparameters and monitoring the average diffRBM units’ score (11) on the validation sets (Figure S3A-B). We also performed additional checks on the diffRBM units’ AUC of immunogenic *vs* non-immunogenic discrimination with different hyperparameters (Figure S3C) and in a control case (Figure S3D).

For the model of T-cell specificity, the background is given by an RBM trained on a random subsample of 10^6^ CDR3*β* sequences either from the bulk repertoires in [Emerson et al., 2017] or the ones in [Dean et al., 2015] (section 5.2.2), choosing the optimal RBM architecture (100 hidden units, 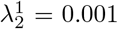) by cross-validation (Figure S14A-B). For cross-validation, we used 50 random partitions of into 10^6^ data for training and 2 × 10^5^ data for validation randomly chosen from the repertoire [Emerson et al., 2017] into training and validation sets (with respectively 80% and 20% of the data). The grid search for optimal hyperparameters for the differential part was then carried out at fixed background model, using the largest dataset of peptide-specific receptors (NLVPMVATV) and partitioning it 50 times at random into training and validation sets (with respectively 80% and 20% of the data, Figure S14C-F).

### 5.4 Alternative approaches

We compared diffRBM performance either to alternative architectures for differential models (diffRBM linear, PWM-based approach, SONIA) or to fully supervised, background-free models (classifiers).

#### 5.4.1 DiffRBM linear

DiffRBM linear is a diffRBM architecture where the differential part is specified by single-site fields 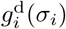 (without the addition of weights 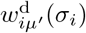).

#### 5.4.2 PWM-based approach

We considered a type of differential model based entirely on the information in single-site amino frequency, where both the background model and the differential part are specified only by single-site fields 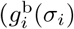 and 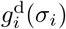 respectively). The equations for learning 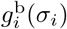 and 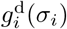 are:

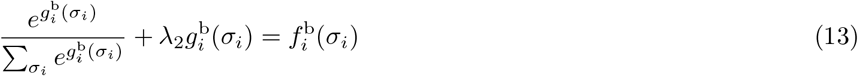

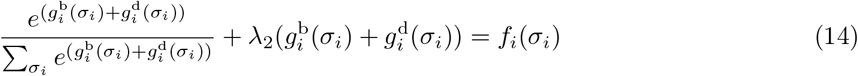

where 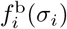 is the frequency of amino acid *σ_i_* at position *i* in the background dataset 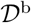, *f_i_*(*σ_i_*) is the frequency of amino acid *σ_i_* at position *i* in the dataset 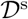 and *i* = 1, …, *N*. We use the same *L*_2_ regularization penalty that we chosen for the RBM fields, and the parameter λ_2_ sets its strength (like for the RBM-based approaches, 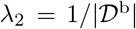 for background models and 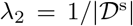 for differential part). This approach is equivalent to learning from the background and selected datasets Position Weight Matrices (PWMs), probabilistic models of amino acid usage that treat all sequence positions as independent. By defining 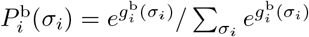 and 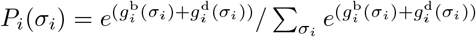 and given the independent-site assumption of PWMs, the background data PWM probability and the selected data PWM probability are respectively recovered as 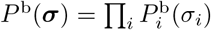 and 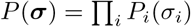. Since 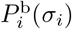 and *P_i_*(*σ_i_*) are learnt to closely reproduce the frequencies 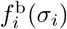 and *f_i_*(*σ_i_*), their predictions are the ones that are inferable from single-site amino acid frequency alone in the respective training datasets (hence in Figures 2, 5, S4, S5, S15 we refer to such predictions by ‘Amino acid frequency’). The prediction by the differential fields can be rewritten as:

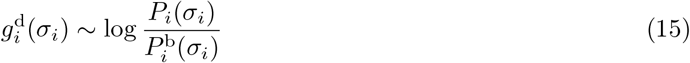

*i.e*., the differential parameters in this case simply corresponds to the log ratio of the PWM learnt on selected data and the PWM learnt on the background data, and measures the site-specific enrichment in amino usage in selected data compared to the background. For a sequence ***σ***, the score based on the differential part is then simply given by the log-likelihood ratio:

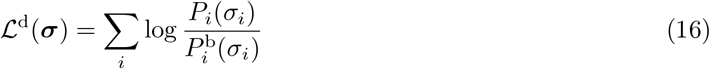

To estimate the entropy of PWM-based approaches (Figure S20C), we used expressions (12) with PWM probabilities for *P*^b^(***σ***) and *P*(***σ***).

#### 5.4.3 SONIA

We implemented a SONIA model using the software package soNNia from [Isacchini et al., 2021] and setting the option *deep=False*. For consistency with the background of diffRBM, we first learn a background model from 10^6^ randomly assembled from the universal donor repertoire from [Emerson et al., 2017], next we learn a model from epitope-specific repertoires and we take, as SONIA score, the difference between the scores assigned by these two models. The *L*_2_-type regularization strength is set to the default value (zero); higher regularizations applied to one of our test sets did not provide visible improvement of the log-likelihood.

#### 5.4.4 Classifier of immunogenic peptides

As a term of comparison for the immunogenicity model, we implemented first a linear (logistic) classifier. (see Model C1 in Table S2), which we trained by minimization of a binary cross-entropy loss. To find an optimal classifier, we also searched among different neural-network architectures trained to discriminate the immunogenic and non-immunogenic data retrieved from IEDB (section 5.2.1). We used the same 50 validation sets held out from the training set of immunogenic, HLA-A*02:01-presented peptides utilized for the model selection of diffRBM. In an analogous way, we randomly partitioned the set of non-immunogenic, HLA-A*02:01-presented peptides into training/test sets (with respectively 80%/20% of the data), repeating the partition 50 times, and we further partitioned the training sets to hold out at each repetition a validation set with 20% of sequences. We trained the models varying the number and width of hidden layers (Table S2), as well as the weight decay. The ‘optimal classifier’ performance for HLA-A*02:01 (Figure 4A) is obtained by selecting for each partition the best performing architecture on the validation set and evaluating its AUC on the test set. Next, we identified the architecture giving the maximal average AUC of discrimination between the immunogenic and non-immunogenic peptides in the validation sets (Model C8 in Table S2, see Figure S11) and we used it to estimate the ‘optimal classifier’ average AUC for the HLA-B*07:02 and HLA-B*35:01 models (Figure 4C-D). Our numerical implementation is based on the PyTorch library. Training is performed in mini-batches of 64 sequences, by the AdamW optimizer with weight decay [Loshchilov and Hutter, 2019], learning rate 0.001, for 500 epochs.

#### 5.4.5 *k*-NN based classifier

Following [Weber et al., 2021], we built a *k*-Nearest Neighbours (*k*-NN) based classifier to distinguish TCRs recognizing a specific antigen from generic ones. It works by computing the Levenshtein distance of the CDR3*β* under analysis with respect to a set of positive examples (training set of antigen-specific CDR3*β*) and to a set of negative examples (training set of bulk repertoire CDR3*β*). In the computation of the Levenshtein distance, the V and J segments are used as well (different V or J segments increase the distance by 1). Then the average distances from the *k* nearest neighbours are computed for both the positive and negative examples, and a score is computed as the difference between the two average distances.

The only parameter of this model, *k*, has been fixed for the largest dataset of positive examples (TCRs reactive to the NLVPMVATV peptide) by cross-validation. We split the positive and negative data in 50 independent training and test sets (with respectively 80% and 20% of the sequences); from each training set, a portion of 20% of its data is held out as validation set. Next *k*, for each partition, is fixed by maximizing the model performance on the validation set, and the model with the best *k* is evaluated on the test set. For the other datasets of positive examples, which are smaller, we used *k* = 26, obtained as the *k* for which the average AUC on the validation set is maximal (Figure S18). The performance of the *k*-NN based classifier was evaluated as described in section 5.5.

#### 5.4.6 NetTCR-2.0

NetTCR-2.0 [Montemurro et al., 2021] is a convolutional neural network method trained from sets of pairs of peptides and cognate TCR sequences (*β* chain only or *α* + *β* chains). To re-train and evaluate it with our datasets in the *β* chain only version, we replaced the sets of CDR3*β* specific to the 4 epitopes of interest (M1_58_, pp65_495_, BMLF1_280_, S_269_) with our VDJdb-derived datasets. We kept all the other NetTCR training data (the 3 peptides different from our 4 peptides of interest with their specific CDR3*β* as well as their control data for these additional peptides). Re-training NetTCR-2.0 with only the 4 epitope-specific sets of CDR3*β* we considered gave AUCs in the same range of values as for the case with 7 peptides. In all cases our training of NetTCR-2.0 uses the same default settings as [Montemurro et al., 2021].

### 5.5 Classification performance

Having found the optimal diffRBM architecture (section 5.3) for the immunogenicity model, for each of the 3 HLA types considered, we trained 50 HLA-specific models on the original training sets (consisting of the 80% of the full datasets available) and we assessed their average performance over the corresponding 50 choices of the test set. In particular, we tested the ability of the HLA-specific immunogenicity models to identify new immunogenic peptides by the Receiver Operating Characteristic curve (ROC). For each of the 50 repetitions, we assigned scores of immunogenicity predicted by a given HLA-specific model (given by the diffRBM units’ score 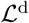 of equation 11) to the sequences of the test set of positives (immunogenic peptides with the HLA type under consideration) and of negatives (a test set of non-immunogenic peptides presented by the same HLA). Varying the threshold score value to discriminate positives from negatives, we obtained the ROC curve describing the fraction of immunogenic peptides predicted by the models’ scores, against the fraction of predicted non-immunogenic ones. We took the Area Under the Curve (AUC) as a metric of the models’ ability to discriminate immunogenic from non-immunogenic peptides. We performed the same validation for all the RBM-based approaches (section 5.1, Figure 6), using their corresponding output scores (equation 9 for the full RBM, 10 for background RBM). The performance of the full RBM obtained through the score (9) is by far and large equivalent to the one of an RBM with the same hyperparameters entirely trained, in one step only, from the selected dataset (Figure S10), confirming that there is a gain in performance with the differential learning strategy only when we focus on the differential units and their parameters. For the diffRBM linear approach (section 5.4), the scores (9) and (11) contain, for the differential part, only fields 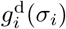; in the PWM-based approach, we used (16) (Figure S7).

We also performed a leave-one-organism-out cross validation, whereby we divided peptides by the organism of origin, we held out as test set only the immunogenic and non-immunogenic peptides from the same organism and trained the models on the peptides from all the other organisms (Figure S9). We considered in the test sets only the organisms for which at least 15 immunogenic and non-immunogenic peptides could be retrieved from IEDB. HLA-A*02:01 is the only allele for which we found sufficient data for this validation.

Also negatives were randomly divided into 50 training and test sets. Training sets of negatives were used to train diffRBM units for non-immunogenic peptides (assigning scores that we will denote as 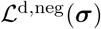 to distinguish it from the scores of immunogenicity 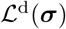) and the classifiers of immunogenicity (section 5.4.4). The classifiers output a probability of being immunogenic which is used as the score of immunogenicity for calculating the AUC on the 50 test sets of positives and negatives. To evaluate the AUC for the approach denoted as ‘diffRBM units (difference)’ (Figures 4, S12), we considered the score given, for each sequence in the test sets ***σ***, by the difference 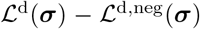. Given that the background model is the same for both the sets of differential units, it can be seen from equations (9) and (11) that the score 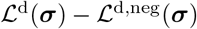 gives the same result as the difference of the full RBM scores 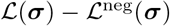 (Figure S10).

We followed the same procedure to train and evaluate the models of T-cell response specificity (Figures 6, S16, S17, S19). In particular, to test the T-cell specificity models’ ability to identify new peptide-specific receptors, we performed an AUC-based assessment of predictive performance analogous to the one described above for peptides, using, as positives, the receptors with the same peptide-specificity from the held-out test sets and, as negatives, a subset of generic receptors from the bulk repertoire (either from [Emerson et al., 2017] or [Dean et al., 2015]), randomly drawn at each repetition with the same size as the positive test set and with no overlap with the 10^6^ sequences of the training set of the background model. All the approaches (diffRBM and alternatives versions, SONIA, *k*-NN, NetTCR-2.0, TCRex) are trained and tested on 50 independent random partitions of both positives and negatives into training and test sets, and the performance shown in Figures 6, S16, S17, S19 is the average AUC over these 50 partitions. As negative set to train the supervised approaches (*k*-NN, NetTCR-2.0) we took, for each training repetition, a subset of the bulk-repertoire dataset from [Emerson et al., 2017] with the same size as the positive training set.

### 5.6 PRIME tool

PRIME [Schmidt et al., 2021] was downloaded from https://github.com/GfellerLab/PRIME. Being it not possible to re-train PRIME on our datasets for a fair comparison of performance, we simply evaluated it on the set of immunogenic and non-immunogenic peptides we collected and we obtained a discrimination performance with AUC=0.53 for peptides specific to HLA-A*02:01, AUC=0.45 for HLA-B*07:02 and AUC=0.56 for HLA-B*35:01. Since we set out to predict immunogenicity conditioned on binding to a given HLA allele, PRIME was run in the mono-allelic mode (*e.g*. with option *-a A0201* for the case of HLA-A*02:01); in general, however, different results are obtained by adding more alleles, where then the best presenting allele according to the predictor is taken.

### 5.7 Contact prediction

#### 5.7.1 Definition of single-site factors

Given the TCR-pMHC structures retrieved from IEDB, we estimated the peptide sites in contact with the HLA and the TCR, as well as the positions of the receptor’s CDR3*β* region in contact with the peptides (section 5.2.4). For these sequences annotated with structural information, we assessed whether differential models can predict contact positions. To this end, we defined single-site factors *T_i_* from the models’ parameters to be evaluated on each sequence ***σ*** as:

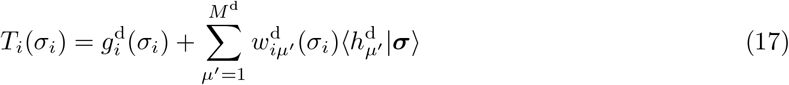

where the average over the differential hidden units 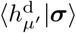 is estimated from a distribution conditional on the sequence ***σ*** that is 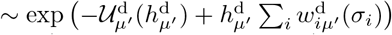. The *T_i_* factors reduce to 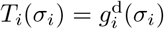 for the diffRBM linear model (Figures S4A-B, S15A-B). From the definition (17), it is clear that singlesite factors measure more generally whether, under a diffRBM model, the amino acid *σ_i_* at position *i*, in the sequence context provided by ***σ***, has high probability to occur among selected sequences, for instance among HLA-specific immunogenic peptides, hence we have used them to quantify residue-specific contributions to immunogenicity (Figure 3).

For the estimation of contacts, the sequence ***σ*** in (17) is represented by either peptides or by receptors’ CDR3*β* sequences when we predict, respectively, the peptide sites in contact with the TCR through the models of immunogenicity (Figure 2C-D) and the CDR3*β* sites in contact with peptides through the models of epitope specificity (Figure 5B-C). Since we interested in a prediction at the level of residues, the models of of epitope specificity used here are defined only on the CDR3*β* amino acid sequence (disregarding the V and J identity). To evaluate contact predictions, for each peptide we use the immunogenicity model corresponding to its HLA type and for each CDR3*β* we use the model corresponding to its epitope specificity. Given the set of *T_i_*(*σ_i_*) from the models for each sequence position *i*, we rank them according to their magnitude and we take the top ranking positions as the model’s prediction on contacts for the sequence ***σ***. In the case of CDR3*β* sequences, we consider only non-gap positions for such ranking.

As a term of comparison, we considered, single-site factors estimated from models which assume all positions independent (section 5.4.2). Here the single-site factors reduce to the log-likelihood ratio:

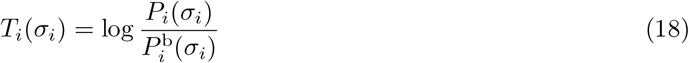

where that *P_i_*(*σ_i_*) and 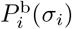 are given by PWMs (section 5.4.2) and hence represent position-specific amino acid frequencies. In the case of the immunogenicity model, the prediction by (18) simply describes the log ratio between the frequency of amino acids in immunogenic peptides relative to the amino acid frequency of all presented peptides from the background dataset (we refer to it as ‘AA frequency ratio (to all presented)’, see Figure 2C-D); in the case of the TCR specificity model (18) gives the log ratio between the frequency of amino acids in peptide-specific CDR3*β* repertoires relative to the amino acid frequency in bulk repertoire (we refer to it as ‘AA frequency ratio (to bulk)’, see Figure 5B-C). In both cases the prediction indicated as ‘Amino acid frequency’ is obtained by estimating and ranking the single-site factors:

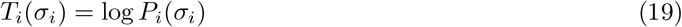

In the case of the immunogenicity model, we have also learnt an independent-site model from non-immunogenic data (that we call *P*^NI^(*σ_i_*)) and we have looked at the single-site factors:

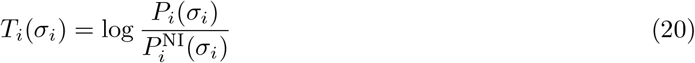

which represents a log ratio of amino acid frequency in immunogenic data relative to non-immunogenic data (this prediction is labelled ‘AA frequency ratio (to non-immunogenic)’ in Figure 2C-D and 3B).

To identify the sites of binding to the HLA, we resort to single-site factors capturing amino acid usage in presented peptides defined as:

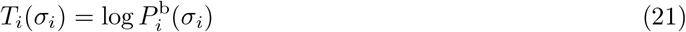

In this case, we look also at the parameters of background RBM, which is learnt to reproduce the amino acid statistics in presented peptides. In particular, consistently with previous analyses [Bravi et al., 2021b], we find that the fields alone identify the anchor sites of binding to the HLA, while the weights tend to capture second-order effects, concentrated at positions 4-8, presumably related to amino acid differences at these positions that might reflect peptide’s conformation in the binding pocket. A case in point was discussed in [Bravi et al., 2021b], where we found that the weights of a single-allele RBM presentation model could capture different binding modes within peptides of the same HLA specificity. Hence we conclude that fields alone are more appropriate for contact prediction and in Figure 2B we use the single-site factors from a fields-only RBM model 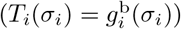 as predictors. In Figure S5A-B we report the comparison to the prediction by a full RBM model 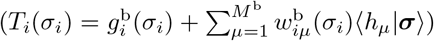, showing only a minimal decrease in performance compared to the fields-only prediction, due to the weights-related effects being mainly second-order ones.

Finally, we have explored an alternative, more general measure of site-specific amino acid importance from the differential units:

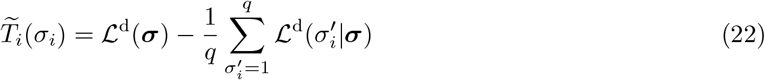

where 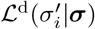 is the differential units’ score of the sequence ***σ*** where position *i* has been mutated from *σ_i_* to 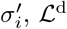 is calculated using equation (11) and *q* is the number of values that 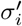 can take. 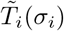 can be hence seen as a measure of the importance of amino acid *σ_i_* at position *i* obtained by comparing its contribution to the likelihood to the one of all other possible *q* amino acids. The diffRBM single-site factors (17) can be derived as a small-weight approximation of the more general definition (22), whose advantage is that *per se* does not depend on the RBM structure of the probability *P*(***σ***). Overall, at the numerical level, the predictions of contacts obtained by ranking sites by (22) are comparable to the ones by (17), see Figures S4A-B, S5A-B, S15A-B.

#### 5.7.2 Prediction assessment via the Positive Predictive Value

Given the models’ predictions of putative contact sites, we assessed their quality by estimating the Positive Predictive Value (Figures 2D, 5C, S4, S5, S15). The Positive Predictive Value for sequence ***σ*** at the ranked position *p* 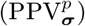 is given by the number of top *p* ranked positions that is included among the contact positions of ***σ*** (true positives), divided by *p* or by number of contacts when this is lower than *p* (all the positives). 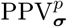 hence hits 1 when *p* is equal to the full length of sequence ***σ***. For a given 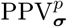, the associated random expectation corresponds to drawing uniformly at random *p* positions and using them to predict the contact positions of ***σ***. The summary values reported in Figures 2D and 5C correspond to the average of 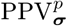 over all the sequences ***σ*** under consideration (respectively, peptides and CDR3*β*) as a function of the number *p* of ranked sequence positions. To correct for the over-representation among the available resolved structures of certain sequences (for example the same peptide and its one-point mutants in contact with different TCRs, or sets of highly similar CDR3*β*s specific to the Influenza peptide GILGFVFTL), we calculate the average PPV^*p*^ at each position *p* as a weighted average:

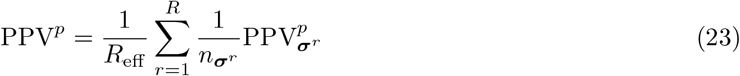

where we reweight the contribution of each sequence to the PPV by a factor 1/*n*_σ_, taking *n_σ_* as the number of sequences that are equal to or mutation away from ***σ***. In (23) we have denoted by *R* the total number of entries under consideration and by *R*_eff_ their effective number obtained as 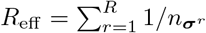. Retaining only unique combinations of sequence and contact positions at the chosen distance cutoffs (4 Å for peptide-TCR contacts, 3.5 Å for peptide-HLA contacts), the number of structures considered for: (*i*) the prediction of contacts with the TCR along the peptide is R = 46 (4 for HLA-B*35:01, 41 for HLA-A*02:01, 1 for HLA-B*07:02); (*ii*) the prediction of contacts with the HLA along the peptide is R =53 (5 for HLA-B*35:01, 47 for HLA-A*02:01, 1 for HLA-B*07:02); (*iii*) the prediction of contacts with the peptide along the CDR3*β* is R =12 (2 for YLQPRTFLL, 3 for NLVPMVATV, 1 for GLCTLVAML, 6 for GILGFVFTL). The corresponding effective numbers are *R*_eff_ = 22.7 for (*i*) and (ii), and *R*_eff_ = 10.3 for (iii). In Figures S4A-B, S5A-B, S15A-B we report the comparison of the average reweighted PPV(equation 23) to the average PPV calculated without reweighting 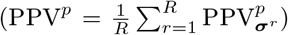, showing that the reweighting does not affect the ranking of performance between different methods.

### 5.8 Mutation costs

The experiments of [Łuksza et al., 2022] on how TCRs cross-react between the NLVPMVATV peptide and its mutants consisted of the following steps: the wild-type peptide (WT) was mutated to every amino acid at every position to obtain 171 mutants (MT); for each MT, its concentration was varied across a 10,000-fold range and the degree activation of 3 WT-specific TCRs was monitored as relative percentage of CD137 expression to determine the TCR cross-reactivity:

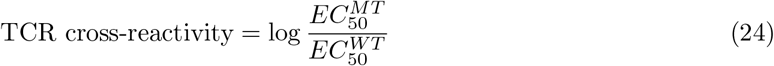

from the WT and MT half maximal effective concentration 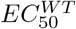 and 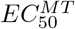 (both measured in *μg/ml*). We took these reported values of TCR cross-reactivity as the experimental mutation costs for each TCR/mutation pair (in the case of what we referred to as ‘non-lethal’ mutations). We defined ‘lethal’ the mutations that were associated to a formally infinite 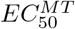 in a given TCR context (*i.e*., whereby TCR response could not be recovered even at the highest concentrations).

For a mutation in sequence ***σ*** at position *i* changing *σ_i_* to 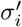, we estimated the model prediction of the mutation cost as:

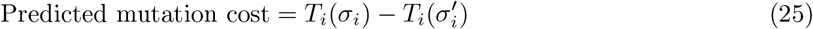

where we took *T_i_* as the single-site factors of (17) for lethal mutations (Figure 3E) and of background RBM (21) for non-lethal mutations (Figure 3G).

To assess whether the distribution towards positive values for lethal mutations is significantly higher than the expectation for generic, non immunogenicity-impacting mutations, we estimated the mutation cost distribution of a ‘control’ case (see Figure 3E) where we drew at random 3000 HLA-A*02:01-presented peptides from the background dataset, and we calculated the costs of all possible amino acid substitutions at each position. The p-value for the difference in these distributions was estimated by the Mann-Whitney U test.

## Acknowledgements

We thank Julien Racle, Nathanael Spisak, Francesco Camaglia, Marta Łuksza, Morten Nielsen and Vadim Karnaukhov for discussions. We are particularly grateful to Giulio Isacchini for help with the data and software. A.D.G. is supported by the European Union’s Horizon 2020 research and innovation programme under the Marie Sklodowska-Curie grant agreement No 101026293. JFdCD acknowledges funding from ANR-19 Decrypted CE30-0021-01 and ANR-17 RBMPro CE30-0021-01. This research was also supported by the European Research Council COG 724208 and ANR-19-CE45-0018 ‘RESP-REP’ from the Agence Nationale de la Recherche.

## Supporting Information

**Figure S1:**
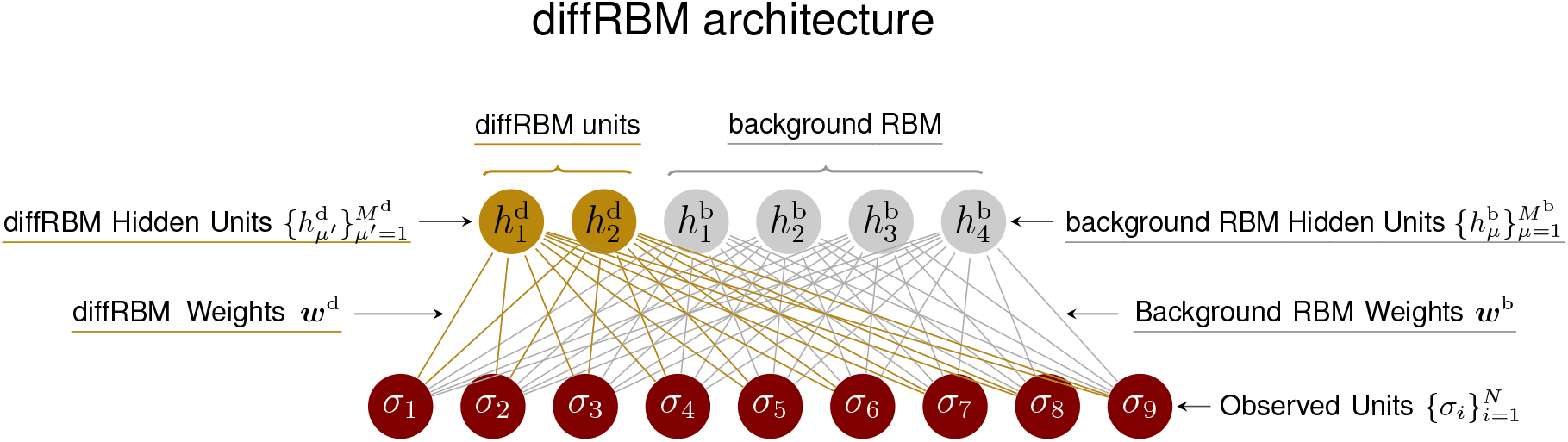
**DiffRBM architecture** recapitulating the mathematical notation used in Methods. In this cartoon example, we have assumed *M*^d^ = 2, *M*^b^ = 4 and *N* = 9 (corresponding to the length of the actual sequence input in the case of peptides, see Figures 2A and S2).

**Figure S2:**
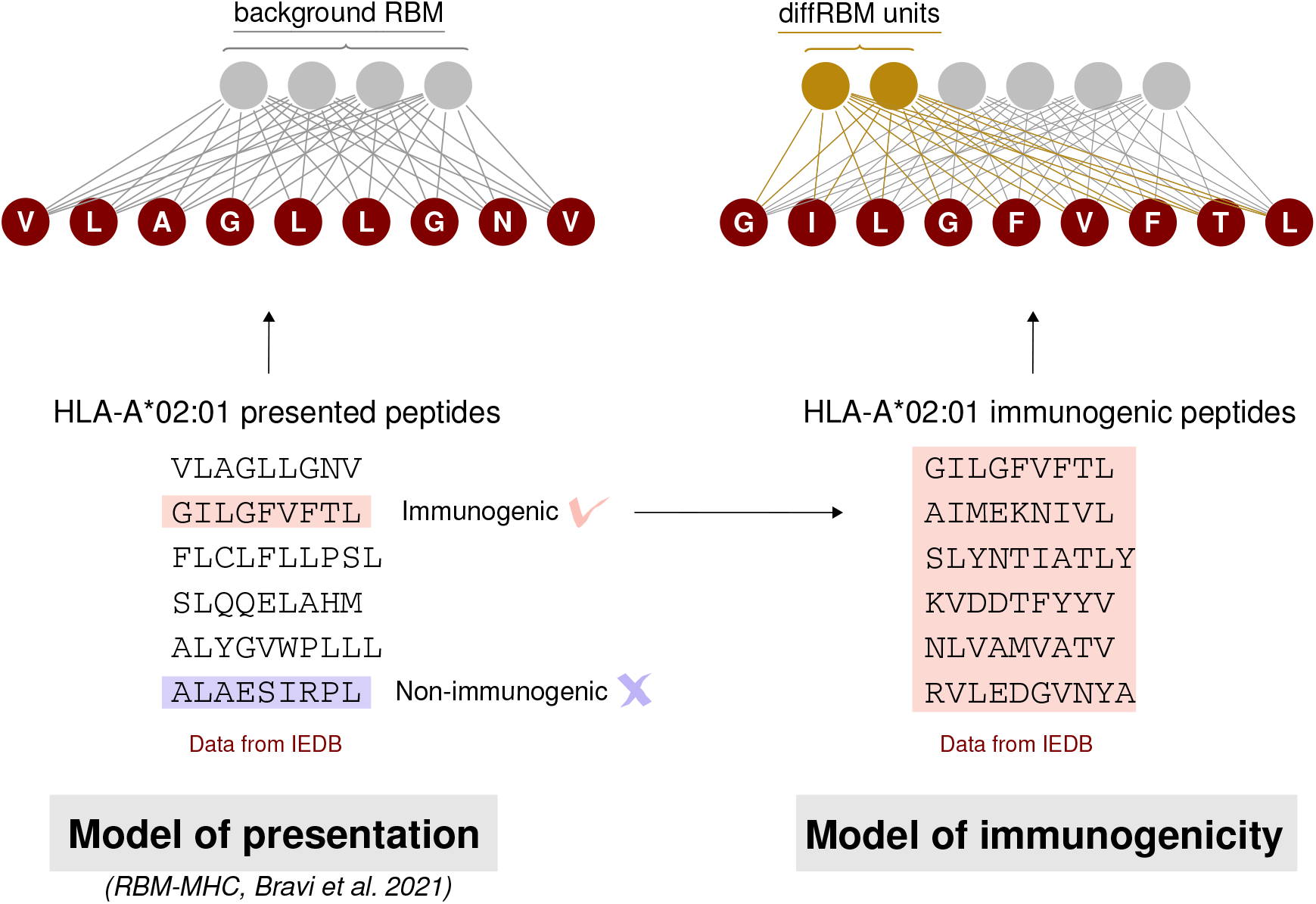
Schematic summary of the construction of a diffRBM model of immunogenicity.

**Figure S3:**
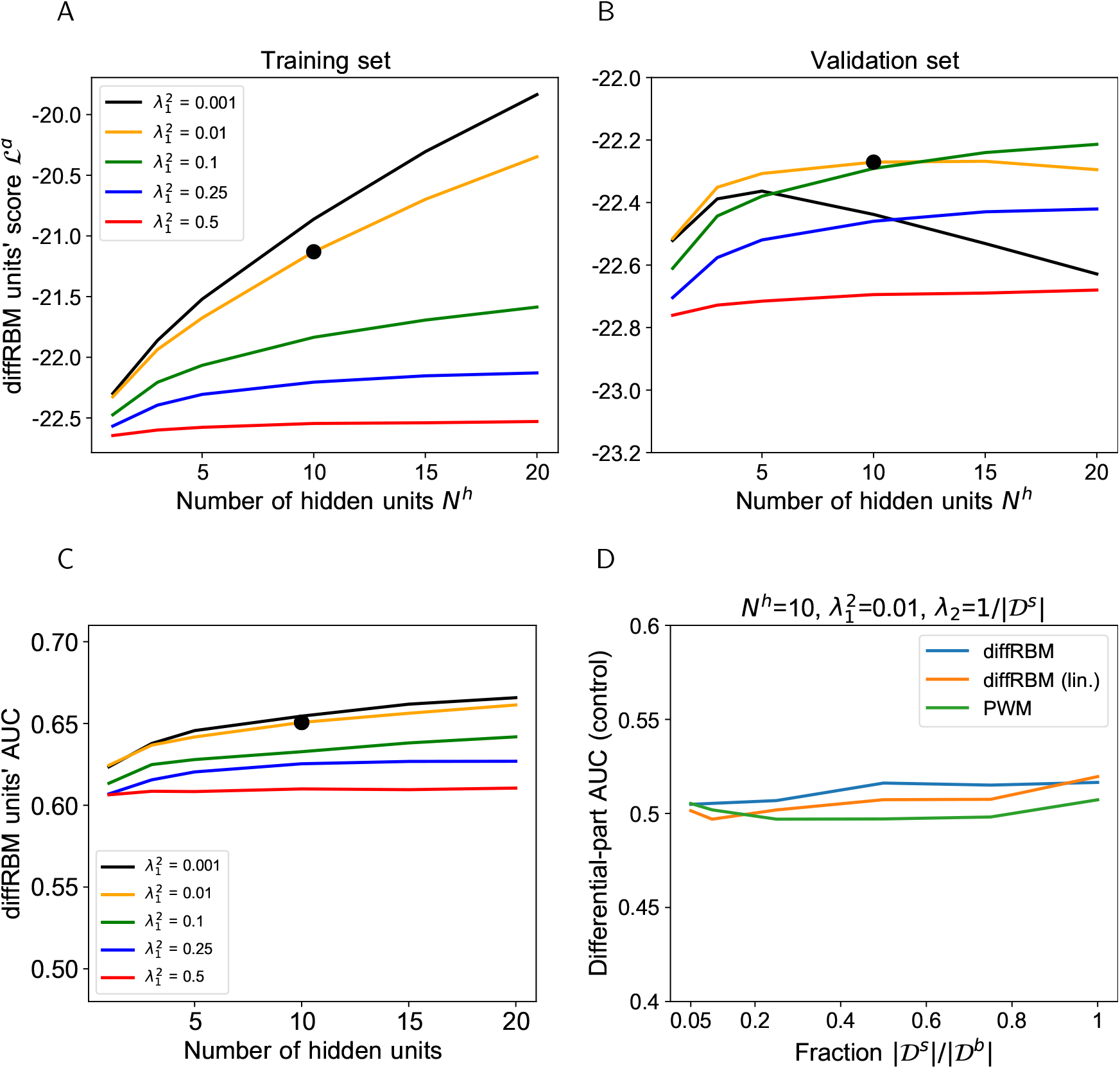
Hyperparametric search for the diffRBM model of immunogenicity. A-B: The hyperparametric search for the diffRBM model of immunogenicity is performed by monitoring the value of the diffRBM units’ score (equation 11) as a function of the number of differential hidden units (*N^h^*) and the regularization 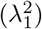. The score values plotted are averages over 50 randomly selected partitions into a training (A) and validation set (B), see Methods. Dataset used: immunogenic peptides presented on HLA-A*02:01 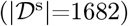. The optimal combination of *N^h^* and 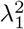 is marked by the black bold dot. C: AUC of discrimination between immunogenic and non-immunogenic in the test set (averaged over 50 random training-test set partitions) given by diffRBMs with different *N*^*h*^ and 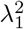 parameters. D: Control of sampling effects on the predictive power of diffRBM with the optimal parameters 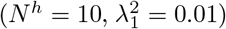, its version where the differential part is linear (diffRBM lin.) and a PWM approach. Here PWM refers to the approaches described in Methods (section 5.4.2), whereby the equivalent of the differential model part is given by the ratio of the PWM learnt on selected data and the PWM learnt on the background data. Differential models are trained on a randomly drawn portion of the background dataset of size 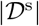, for varying 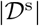, instead of a training set of immunogenic-only sequences, and then the differential part is used to estimate the AUC of discrimination between immunogenic and non-immunogenic sequences in the test set. Differential models, as expected in this control dataset, have no power at discriminating immunogenic from non-immunogenic peptides (AUC ~ 0.5) for all values of the ratio 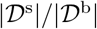.

Table S1: List of TCR-pMHC structures from PDB and estimated contact positions (provided as a separate .xlxs table).

**Figure S4:**
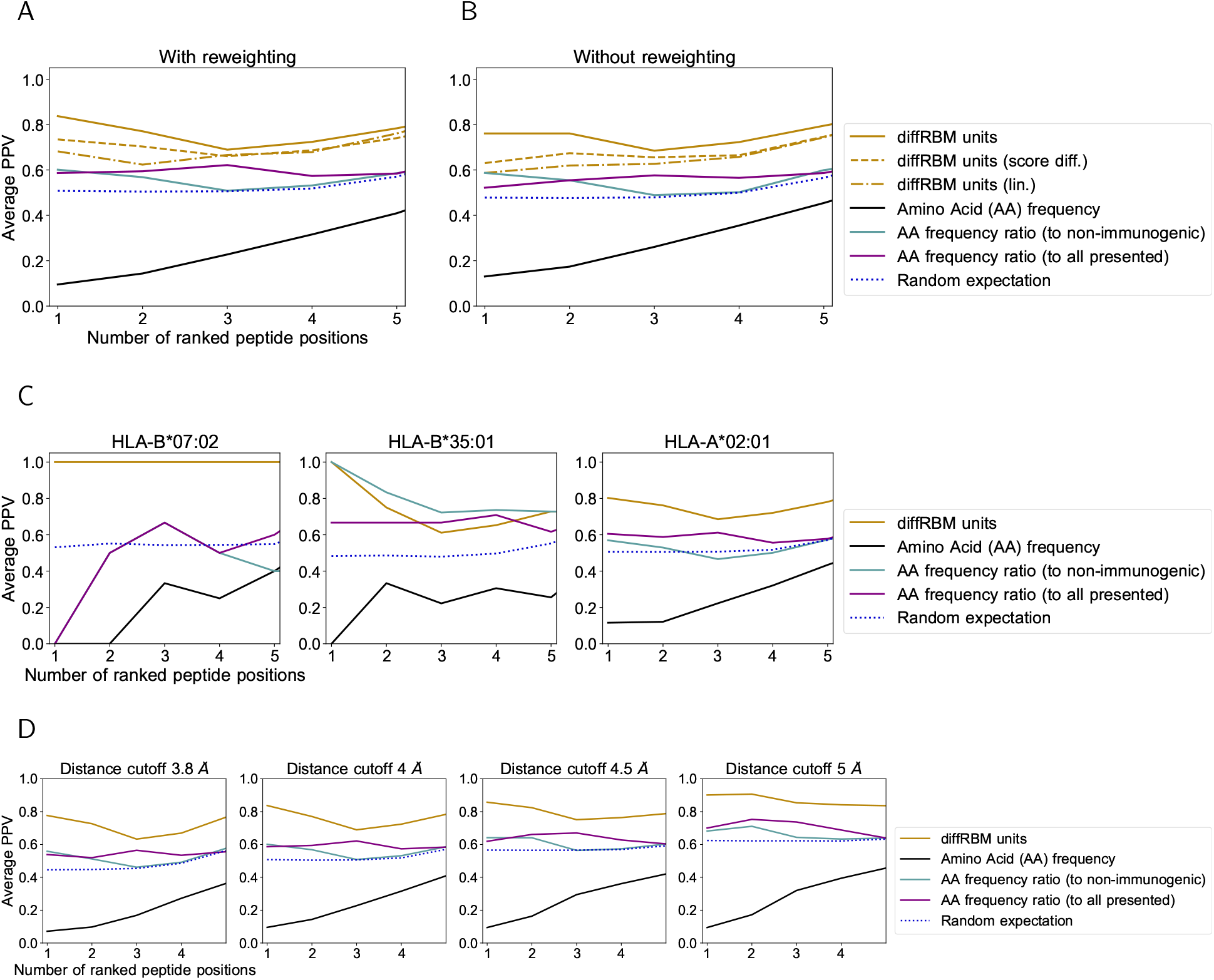
Prediction of peptide contact positions with the TCR. A: The average PPV shown in Figure 2D is compared to the PPV obtained from: the single-site factors from diffRBM units with fields only (‘diffRBM units (lin.)’, dashed-dotted line); a prediction based on a difference of diffRBM units’ scores (equation 22). The later is marked as ‘diffRBM units (score diff.)’ (dashed line) to distinguish it from the prediction by diffRBM single-site factors (17) (simply denoted as ‘diffRBM units’). B is the same plot as A but obtained without applying the reweighting to account for peptide sequence similarity (Methods). C: Same plot as Figure 2D, where predictions are divided by HLA allele. The number of structures considered is as follows: 4 structures for HLA-B*35:01, 41 for HLA-A*02:01, and 1 for HLA-B*07:02. D: Same plot as Figure 2D varying the threshold in Å to define a contact. In all plots, the blue dotted line marks the result of predicting at random the contact positions (Methods).

**Figure S5:**
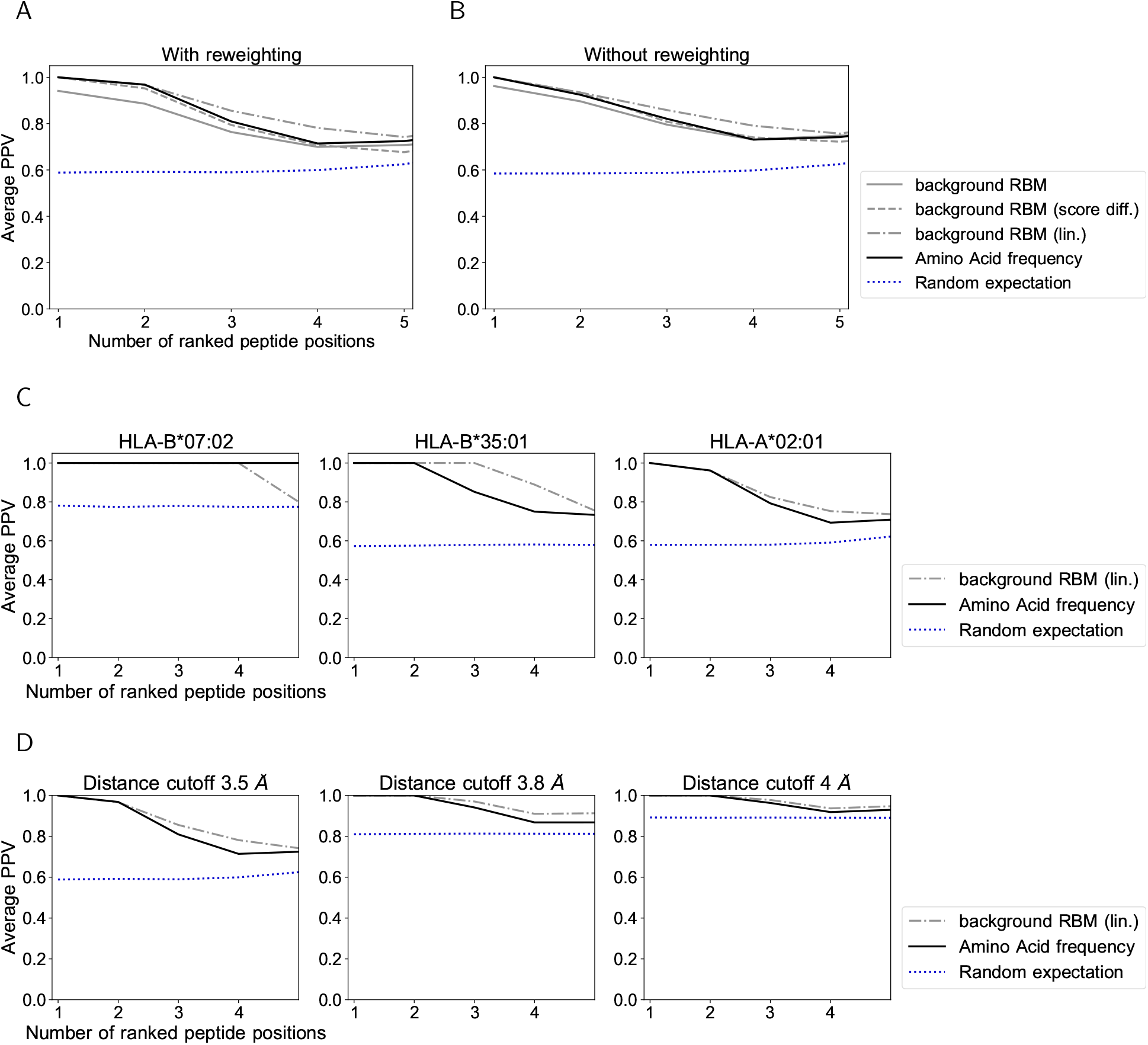
Prediction of peptide contact positions with the HLA. A: The average PPV of contact prediction shown in Figure 2B (inset) is compared to the PPV obtained from: the single-site factors from a background RBM with all parameters (bold gray line); a prediction based on a difference of background RBM log-likelihoods (‘background RBM (score diff.)’, dashed line). B is the same plot as A but obtained without applying the reweighting to account for peptide sequence similarity (Methods). C: Same plot as Figure 2B (inset) where the prediction is shown separately for each HLA (4 structures for HLA-B*35:01, 41 for HLA-A*02:01, and 1 for HLA-B*07:02). D: Same plot as Figure 2B (inset) varying the threshold in Å to define a contact. In all plots, the blue dotted line marks the result of predicting at random the contact positions (Methods).

**Figure S6:**
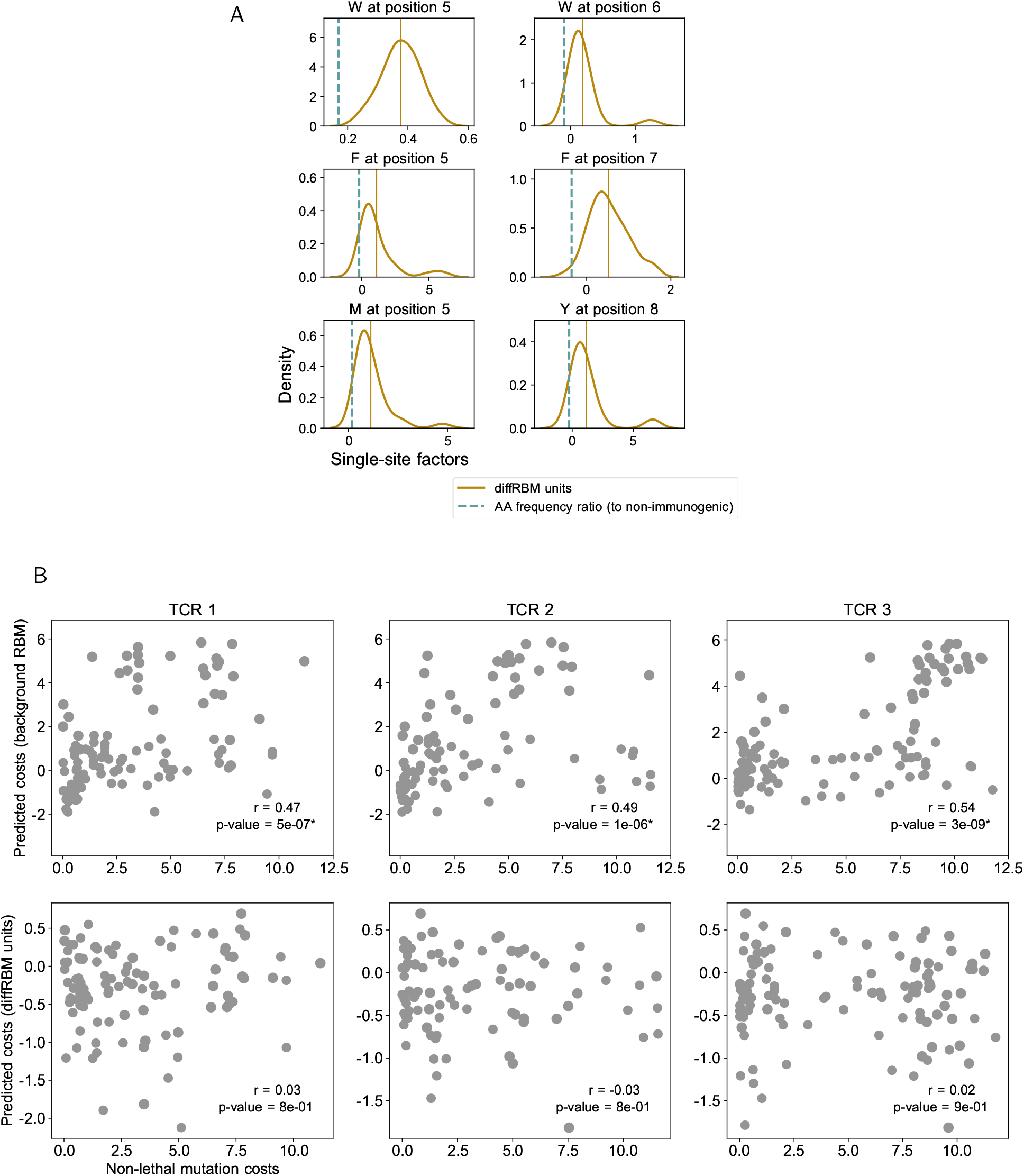
Prediction of immunogenicity-related residues and mutation costs. A: Same as Figure 3B for all the key residues: W at position 5 (39 sequences) and position 6 (20 sequences), F at position 5 (104 sequences) and position 7 (107 sequences); M at position 5 (33 sequences); Y at position 8 (90 sequences). B: Experimental *vs* predicted mutation costs of non-lethal mutations (like in Figure 3G) for all 3 TCRs analyzed in [Łuksza et al., 2022], showing the predictions both by background RBM and the diffRBM units. The Spearmann correlation coefficient *r* for the predictions by background RBM is significant across the 3 TCRs (p-value ≤ 10^−6^).

**Figure S7:**
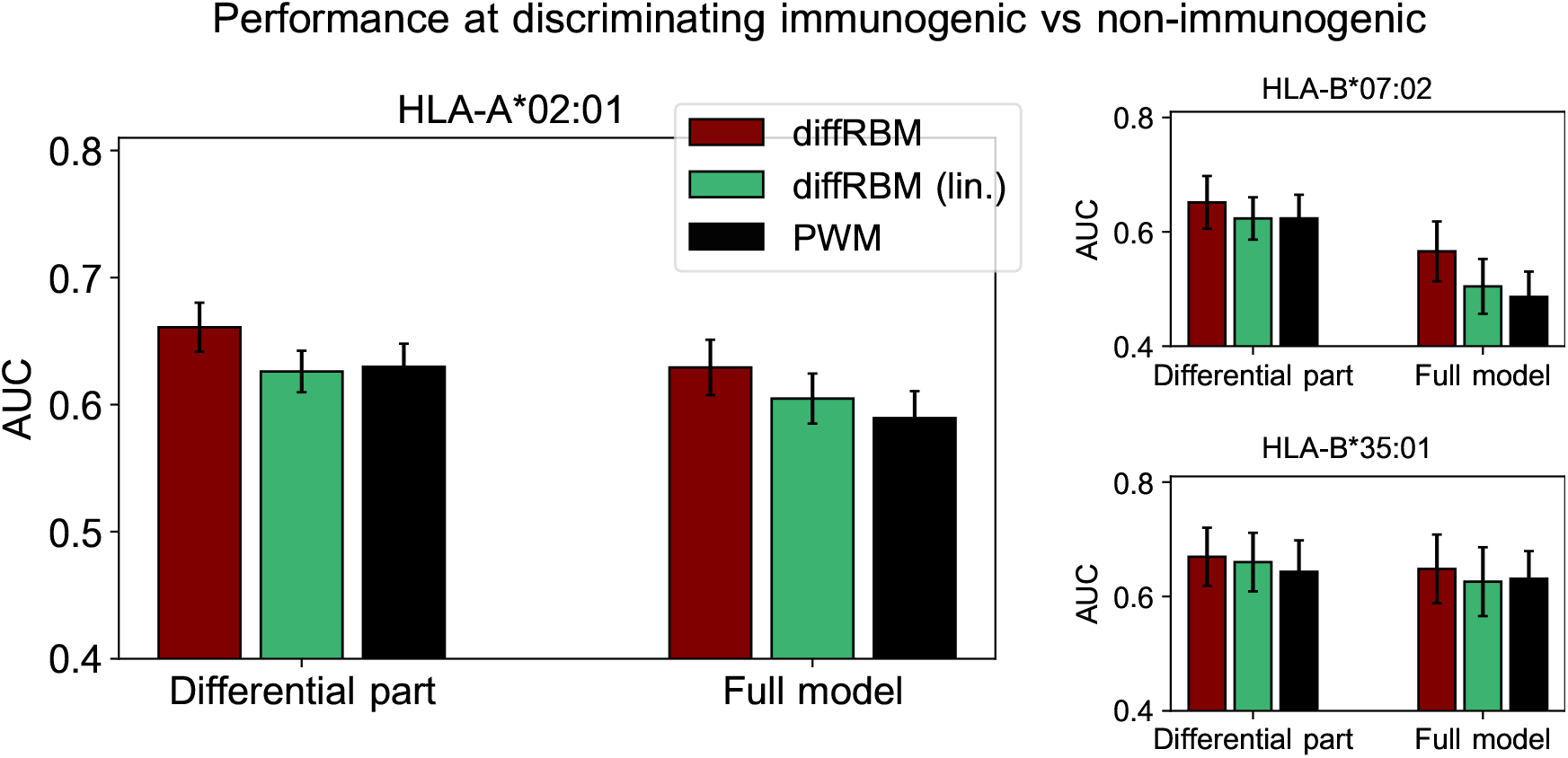
Comparison of performance of differential models of immunogenicity. Comparison of performance, in terms of the AUC of immunogenic *vs* non-immunogenic discrimination, between diffRBM, its version where the differential part is linear (diffRBM lin.) and a PWM approach. Here PWM refers to the approaches described in Methods (section 5.4.2), where the ‘Full model’ is given by the PWM learnt on the selected data (immunogenic peptides) while the ‘Differential part’ corresponds to the ratio of the PWM learnt on selected data and the PWM learnt on the background data. AUCs are averages over 50 training-test partitions and error bars give the corresponding standard deviation (Methods).

**Figure S8:**
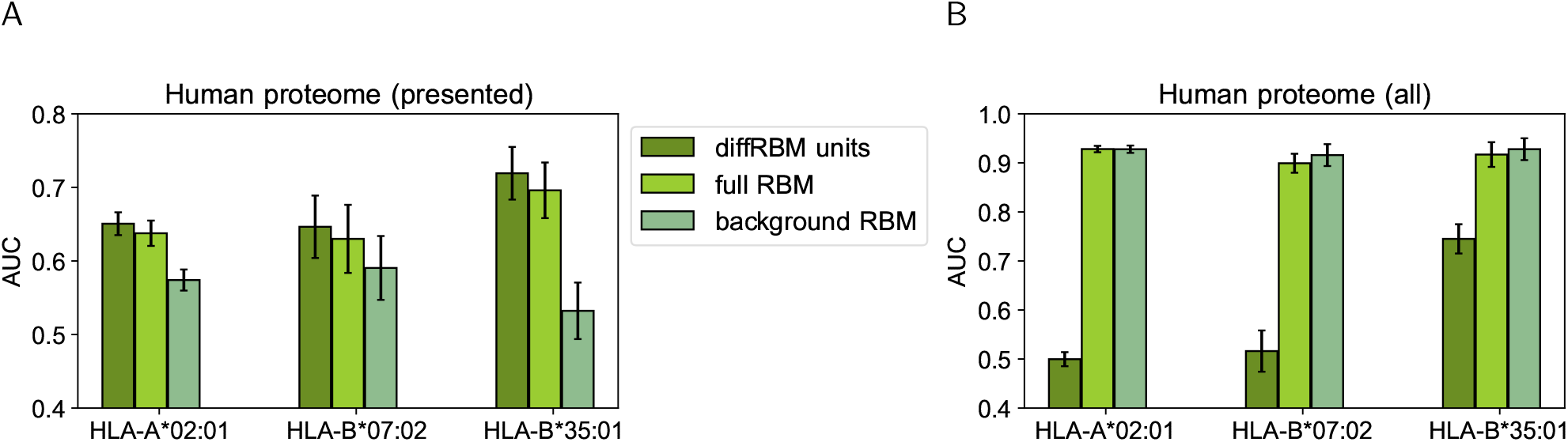
Score comparison between immunogenic peptides and peptides from the human proteome. We have drawn at random 10^5^ peptides from all the possible 9-mers in the human proteome and we have assigned to them scores under the HLA-specific models listed in the legend. We have measured via the AUC the extent to which scores assigned by these models to HLA-specific immunogenic peptides are higher than the ones assigned either to all the 10^5^ human proteome peptides (A) or only to the ones among them predicted to be presented on the corresponding HLA by NetMHCpan4.1 (B). AUCs are averages over 50 training-test partitions and error bars give the corresponding standard deviation (Methods).

**Figure S9:**
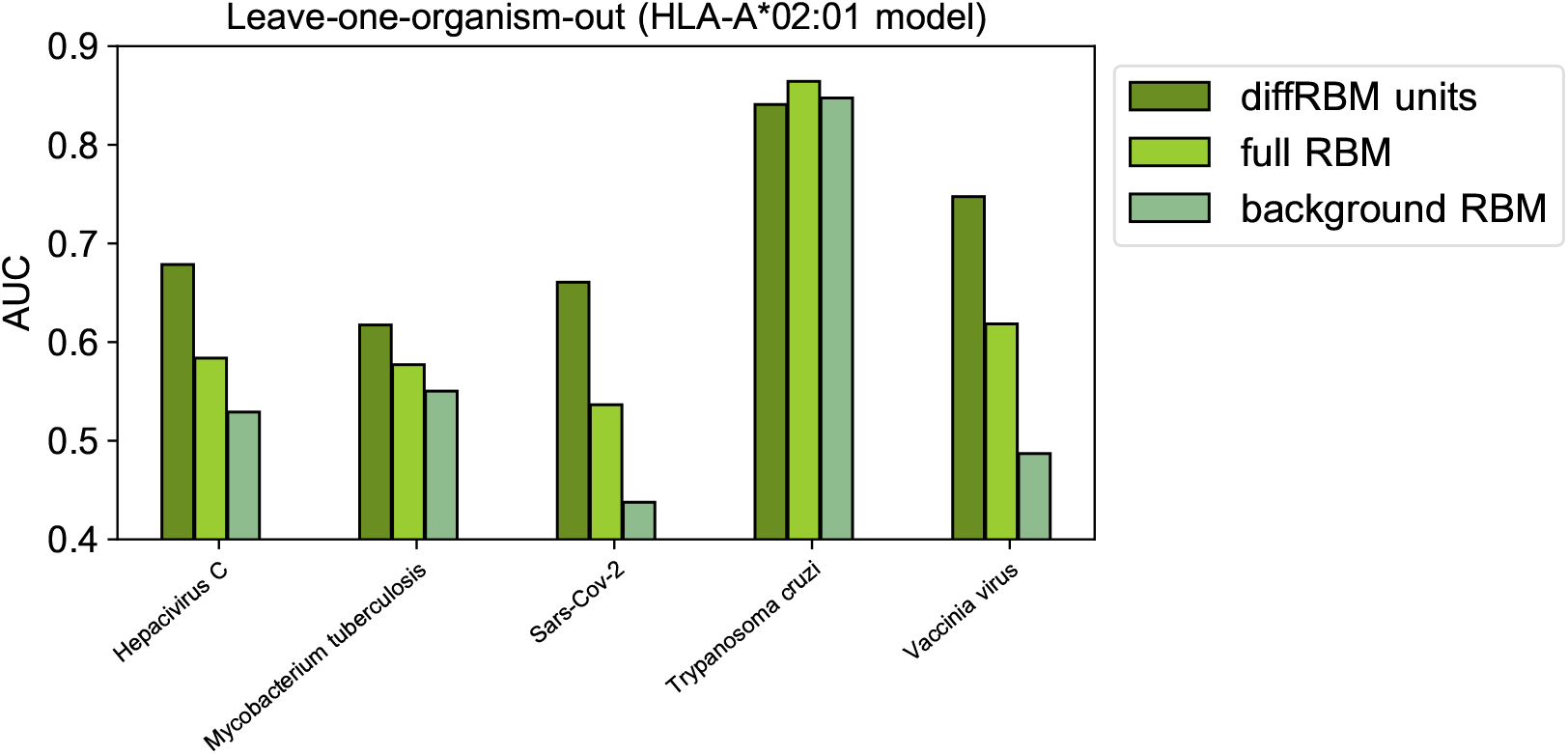
Leave-one-organism-out cross-validation for HLA-A*02:01-specific model (Methods). The case of *Trypanosoma cruzi* visibly constitutes an outlier, whereby the binding affinity to the HLA alone discriminates accurately immunogenic and non immunogenic antigens (AUC=0.85). This result is confirmed by scoring the peptides via NetMHCpan4.1 (AUC=0.85). Excluding this case, the average diffRBM units AUC=0.68, which is comparable to the AUC obtained via standard validation (average AUC=0.66, Figure 4B).

**Figure S10:**
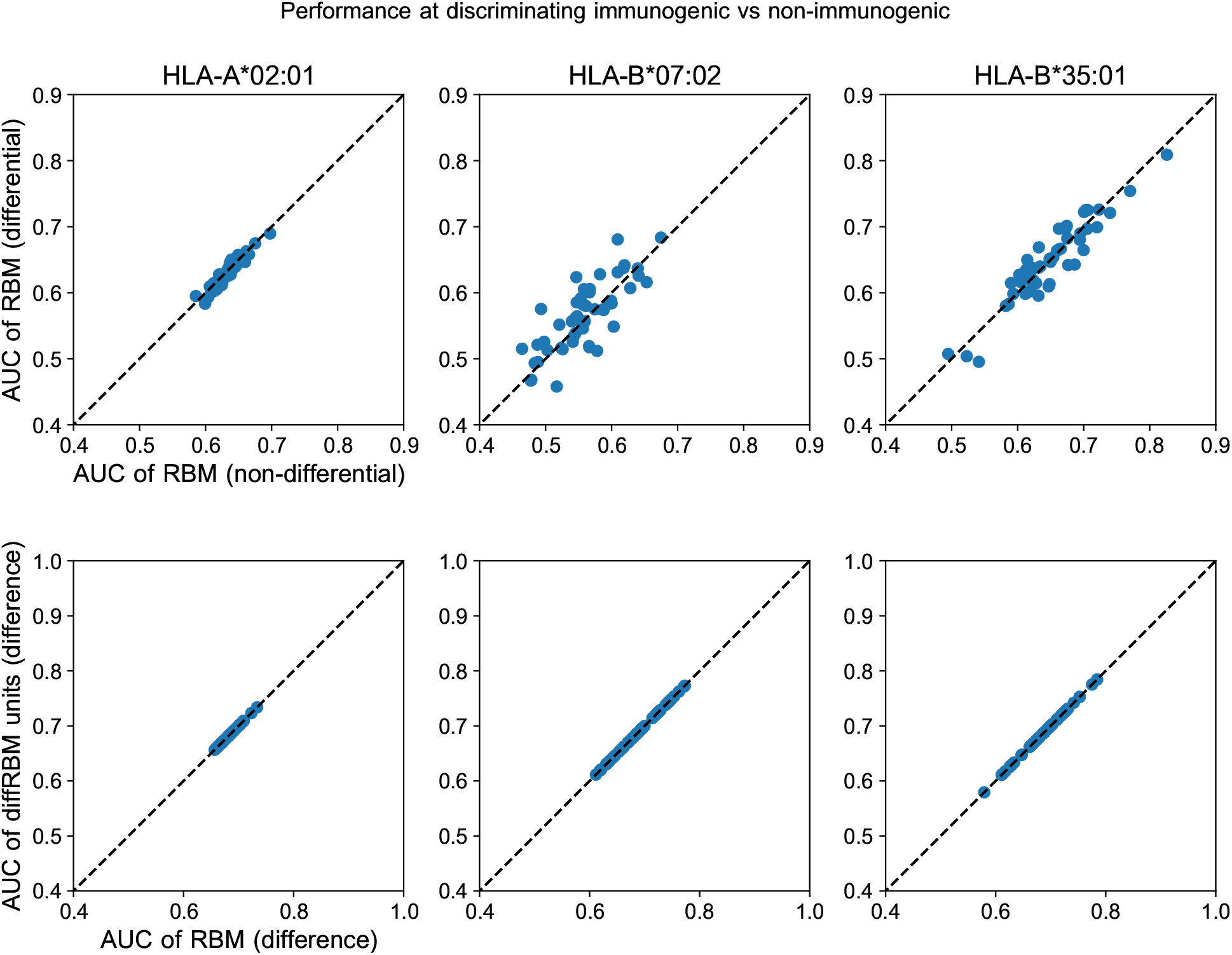
Further comparison of diffRBM and RBM scores. Upper row: the AUC of classification of immunogenic *vs* non-immunogenic peptides given by the full RBM (trained in part on the background dataset and in part on the selected data and indicated as ‘RBM (differential)’) is essentially the same as the one of an RBM with the same hyperparameters trained only on selected data (denoted as ‘RBM (non-differential)’). The comparison is made separately for each HLA. In these scatter plots every point gives the AUC of discrimination of the two models compared for one of the 50 random partitions into training and test sets of the corresponding HLA-specific dataset (Methods). Bottom row: we consider the AUC of immunogenic *vs* non-immunogenic classification obtained through the difference of scores between the diffRBM units trained on immunogenic peptides and the ones trained on non-immunogenic peptides. The scatter plots show that such performance (on the *y*-axis), as it should, is equivalent to the one attained by the difference of the full RBM log-likelihoods (*x*-axis).

**Figure S11:**
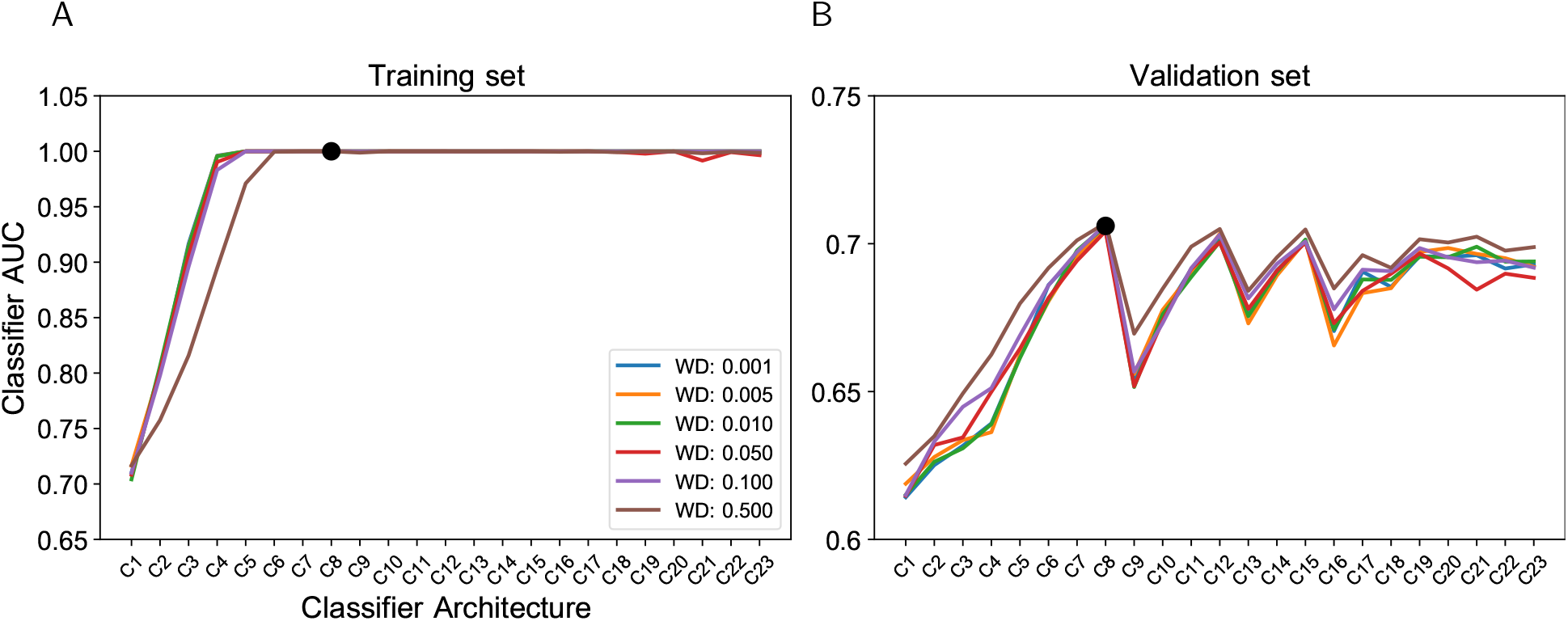
Hyperparametric search for the classifier of immunogenicity. The hyperparametric search for the classifier is based on the optimal performance on the validation dataset at discriminating immunogenic *vs* non-immunogenic peptides as measured by the AUC. We show the mean AUC over 50 random partitions into training set (A) and validation set (B). The labels on the *x*-axis denote different classifier architectures, which are described in Table S2.

**Table S2:** Tested architectures for the classifier of immunogenicity. All classifiers alternate dense linear layers with LeakyReLU [Maas et al., 2013] activation functions. The models differ in the number of layers (column *Depth*) and the width of intermediate layers (column *Intermediate widths*). The first layer is always of input size 180 = 21 × 9, and corresponds to one-hot encoded sequences of length 9 with 20 amino-acid + 1 gap letters. The last layer always has an output of size 1 giving the predicted log-odds that the input sequence is immunogenic. Model C1 has no intermediate layers and corresponds to the linear classifier whose results are shown in Figure 4.

| Id. | Depth | Input | Intermediate widths | Output |
| --- | --- | --- | --- | --- |
| C1 | 2 | 180 | - | 1 |
| C2 | 3 | 180 | 2 | 1 |
| C3 | 3 | 180 | 4 | 1 |
| C4 | 3 | 180 | 8 | 1 |
| C5 | 3 | 180 | 16 | 1 |
| C6 | 3 | 180 | 32 | 1 |
| C7 | 3 | 180 | 64 | 1 |
| C8 | 3 | 180 | 128 | 1 |
| C9 | 4 | 180 | 16, 8 | 1 |
| C10 | 4 | 180 | 32, 8 | 1 |
| C11 | 4 | 180 | 64, 8 | 1 |
| C12 | 4 | 180 | 128, 8 | 1 |
| C13 | 4 | 180 | 32, 16 | 1 |
| C14 | 4 | 180 | 64, 16 | 1 |
| C15 | 4 | 180 | 128, 16 | 1 |
| C16 | 5 | 180 | 32, 16, 8 | 1 |
| C17 | 5 | 180 | 64, 16, 8 | 1 |
| C18 | 5 | 180 | 64, 32, 16 | 1 |
| C19 | 5 | 180 | 128, 32, 16 | 1 |
| C20 | 5 | 180 | 128, 64, 16 | 1 |
| C21 | 5 | 180 | 128, 64, 32 | 1 |
| C22 | 6 | 180 | 128, 64, 32, 8 | 1 |
| C23 | 6 | 180 | 128, 64, 32, 16 | 1 |

**Figure S12:**
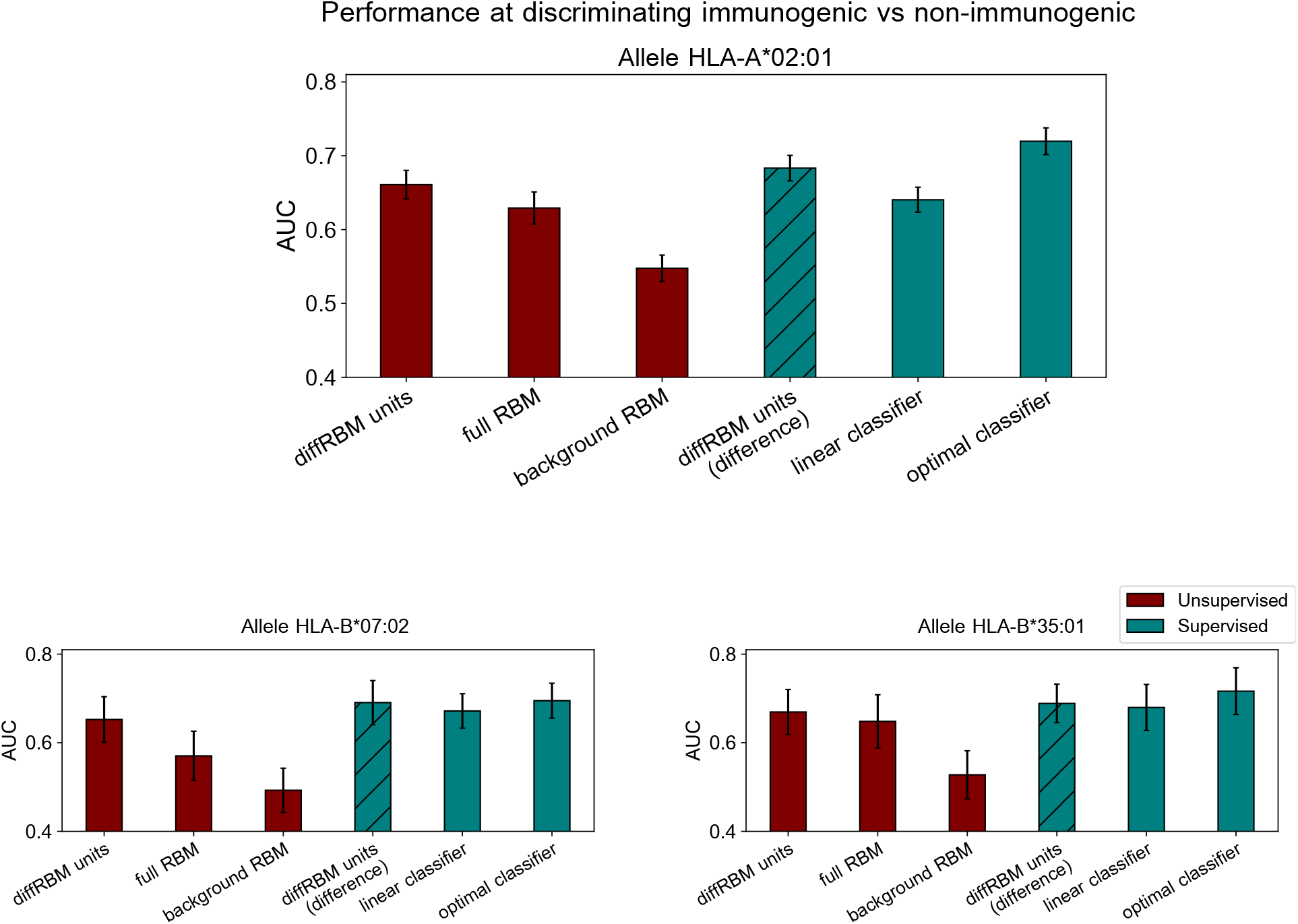
Performance of differential models of immunogenicity with sample reweighting. Performance at discriminating immunogenic *vs* non-immunogenic peptides (like in Figure 4) where a reweighting scheme based on sequence similarity is applied during training (Methods).

**Figure S13:**
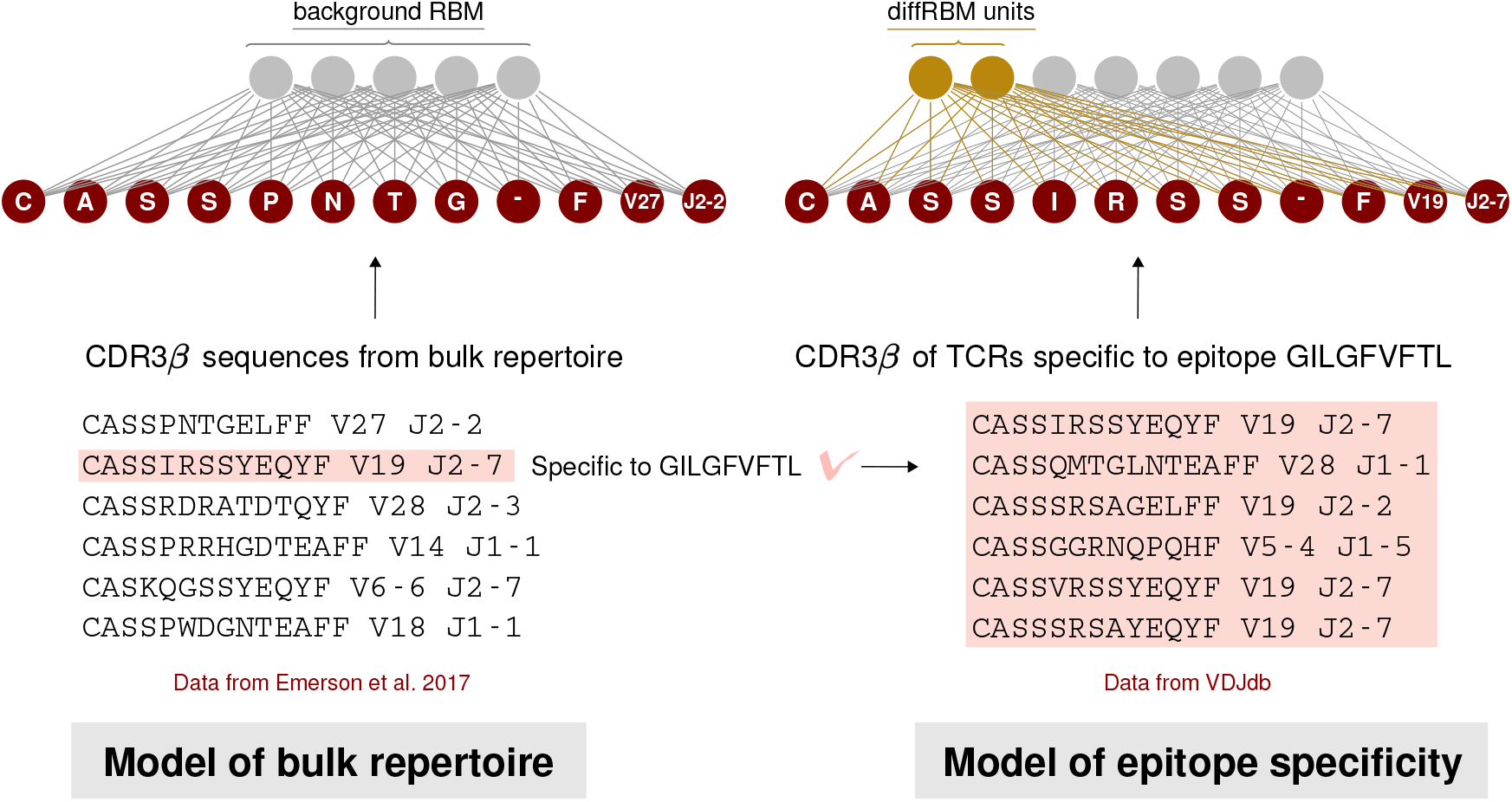
Schematic summary of the construction of a diffRBM model of TCR epitope-specificity.

**Figure S14:**
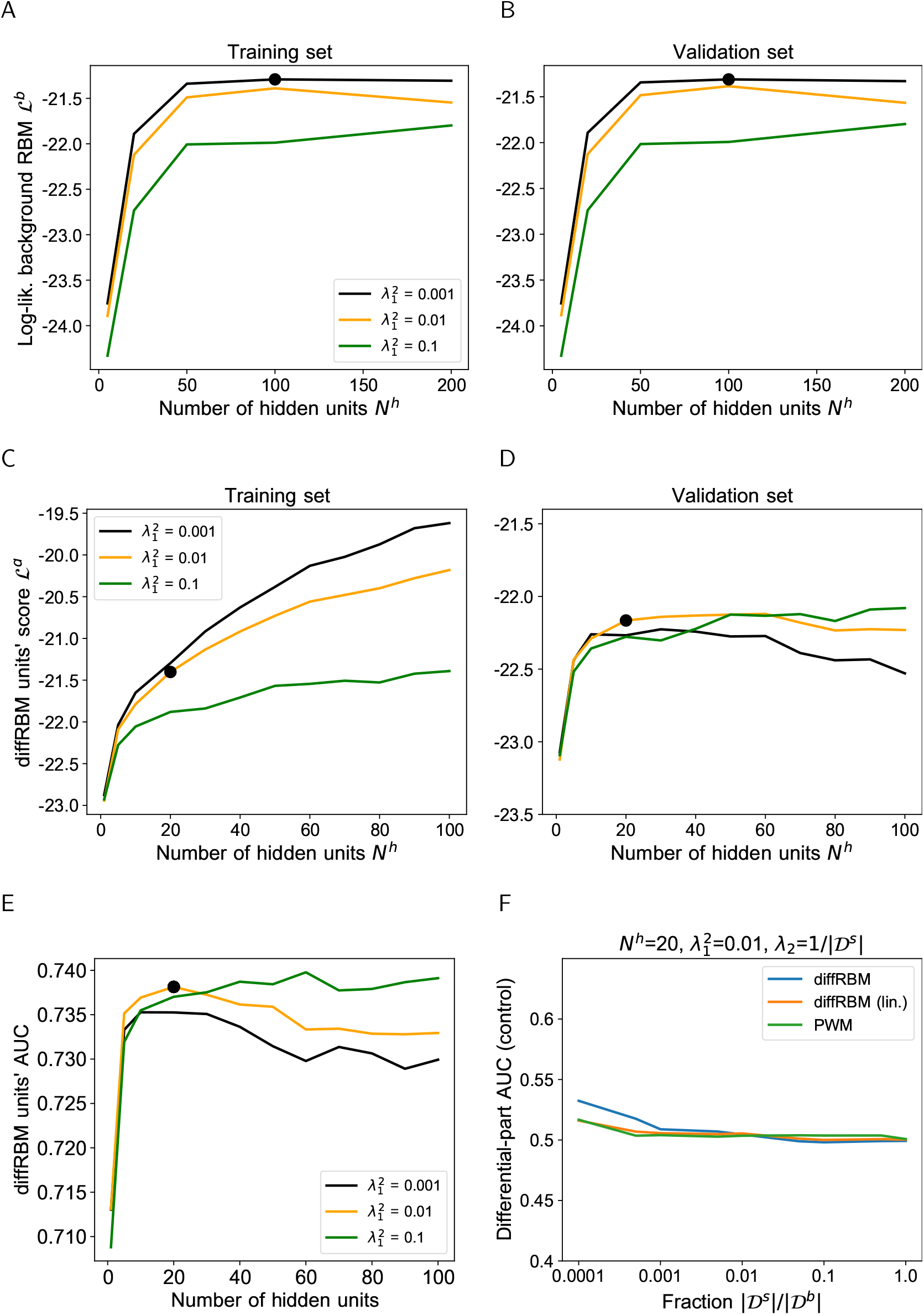
Hyperparametric search for the diffRBM model of TCR specificity. A-B: Hyperparametric search for the background RBM model using the dataset from [Emerson et al., 2017] (Methods). The black bold dot in A-B and C-E marks the combination of *N*^*h*^ and 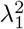 chosen, respectively, for background RBM (*N*^*h*^= 100, 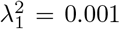) and diffRBM units (*N*^*h*^= 20, 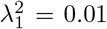). The hyperparametric search for diffRBM is performed in the same way as for the immunogenicity model (Figure S3A-B). The diffRBM units’ scores plotted are averages over 50 random partitions into a training (C) and validation set (D), see Methods. Dataset used: set of sequences specific to the CMV peptide NLVPMVATV 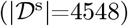. E: AUC of discrimination between epitope-specific and generic sequences in the test set (averaged over 50 random training-test set partitions) varying *N*^*h*^ and 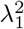. F: As a control, differential models are trained on a randomly drawn portion of the background dataset of size 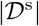, for varying 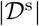, instead of a training set of epitope-specific sequences, and then their differential part is used to estimate the AUC of discrimination between epitope-specific and generic sequences in the test set, see the caption of Figure S3D.

**Figure S15:**
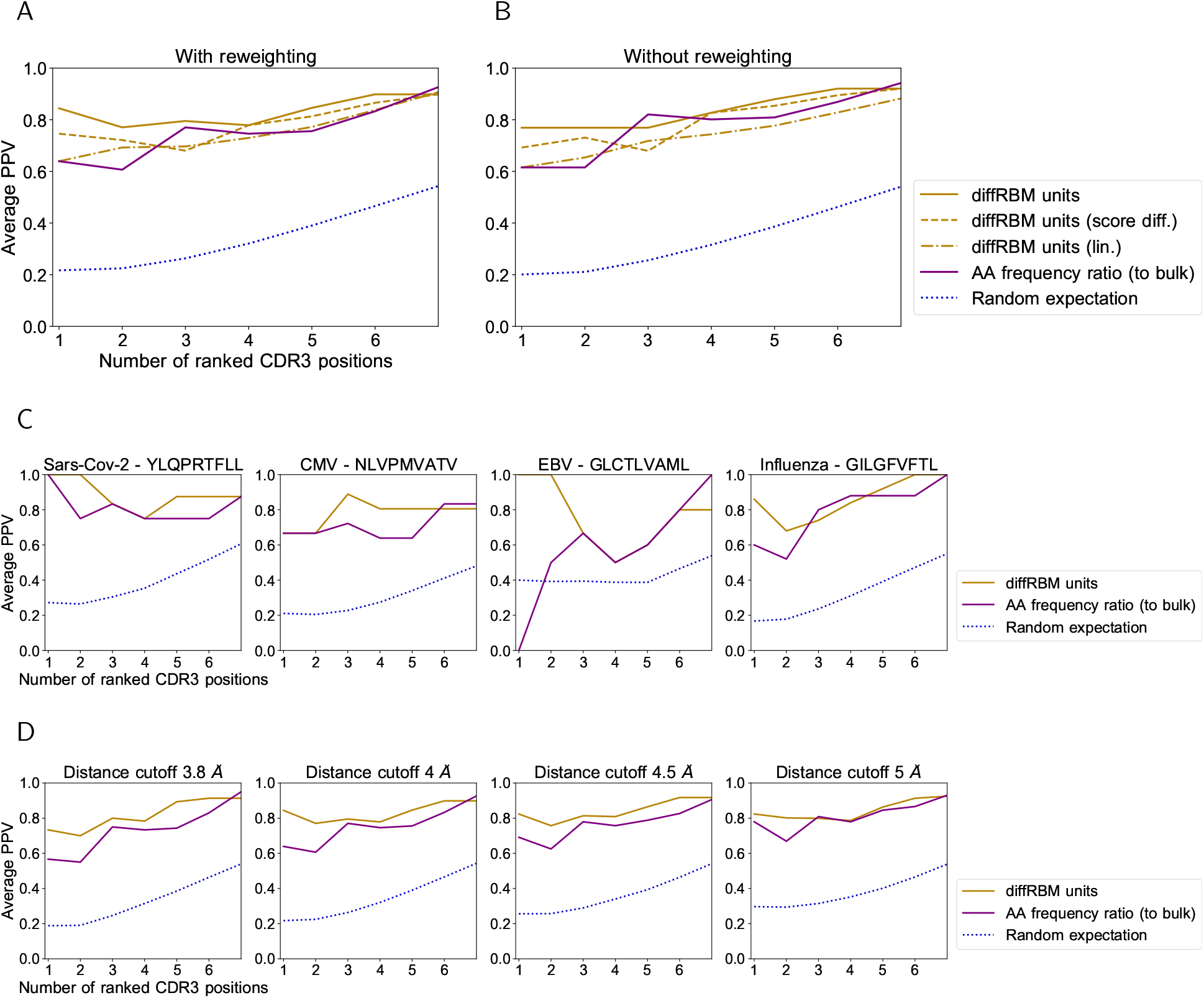
Prediction of CDR3*β* contact positions with the peptide. A: Similarly to Figure S4A, the average PPV of contact prediction shown in Figure 5C is compared to the PPV by two alternative predictors, the expression (22) (‘diffRBM units (score diff.)’, dashed line) and the single-site factors from diffRBM units with fields only (‘diffRBM units (lin.)’, dashed-dotted line). B is the same plot as A but obtained without applying the reweighting to account for peptide sequence similarity (Methods). C: Same plot as Figure 5C, where predictions are divided by epitope specificity. The number of structures considered is as follows: 2 structures for YLQPRTFLL, 3 for NLVPMVATV, 1 for GLCTLVAML, and 6 for GILGFVFTL. D: Same plot as Figure 5C varying the threshold in Å to define a contact. The cutoff 5 Å is the one used in [Calis et al., 2012, Glanville et al., 2017, Ostmeyer et al., 2019, Milighetti et al., 2021]. In all plots, the blue dotted line marks the result of predicting at random the contact positions (Methods).

**Figure S16:**
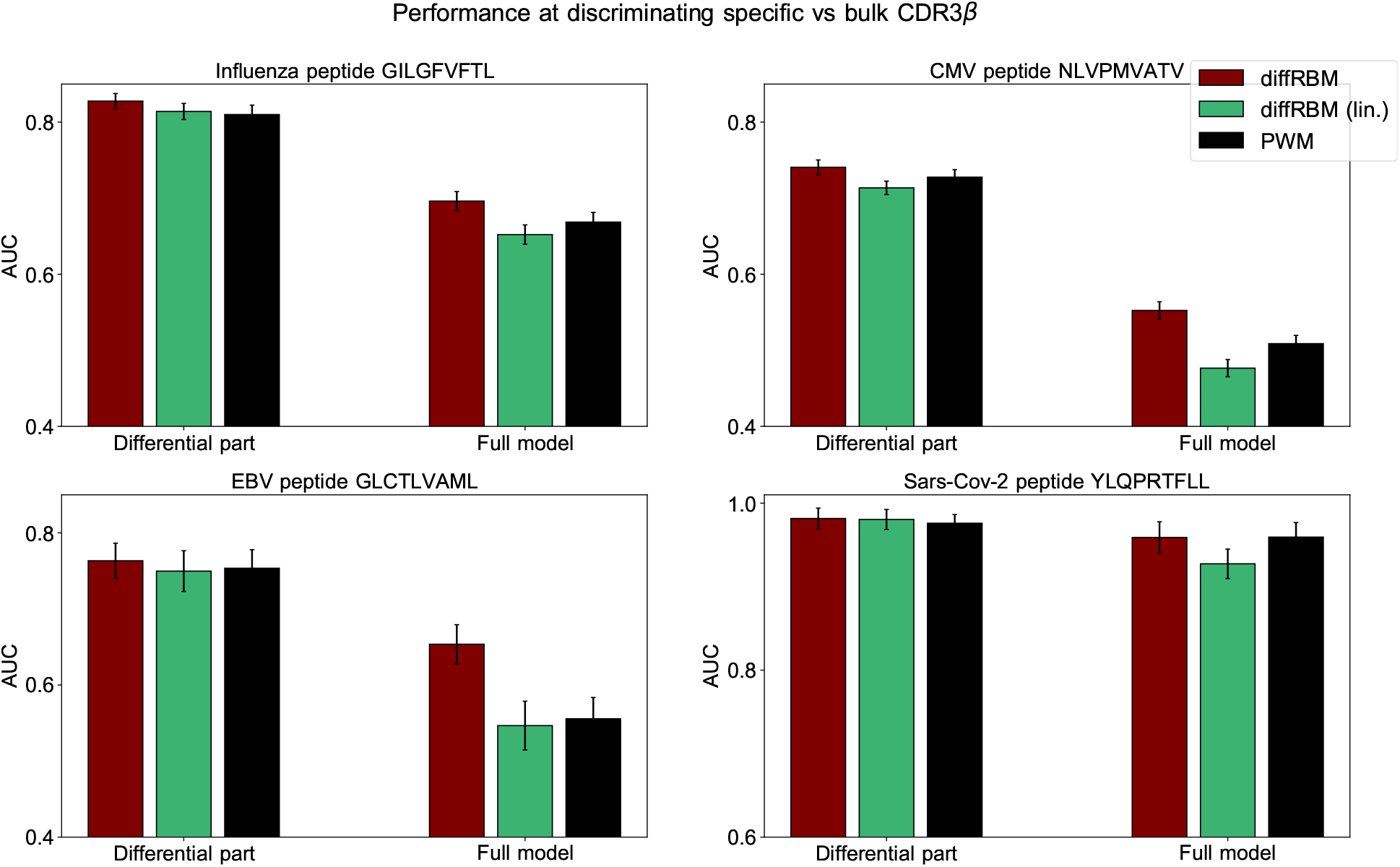
Comparison of performance of differential models of TCR specificity. Comparison of performance, in terms of the AUC of epitope-specific *vs* bulk sequences discrimination, between diffRBM, its version where the differential part is linear (diffRBM lin.) and a PWM approach, see caption of Figure S7.

**Figure S17:**
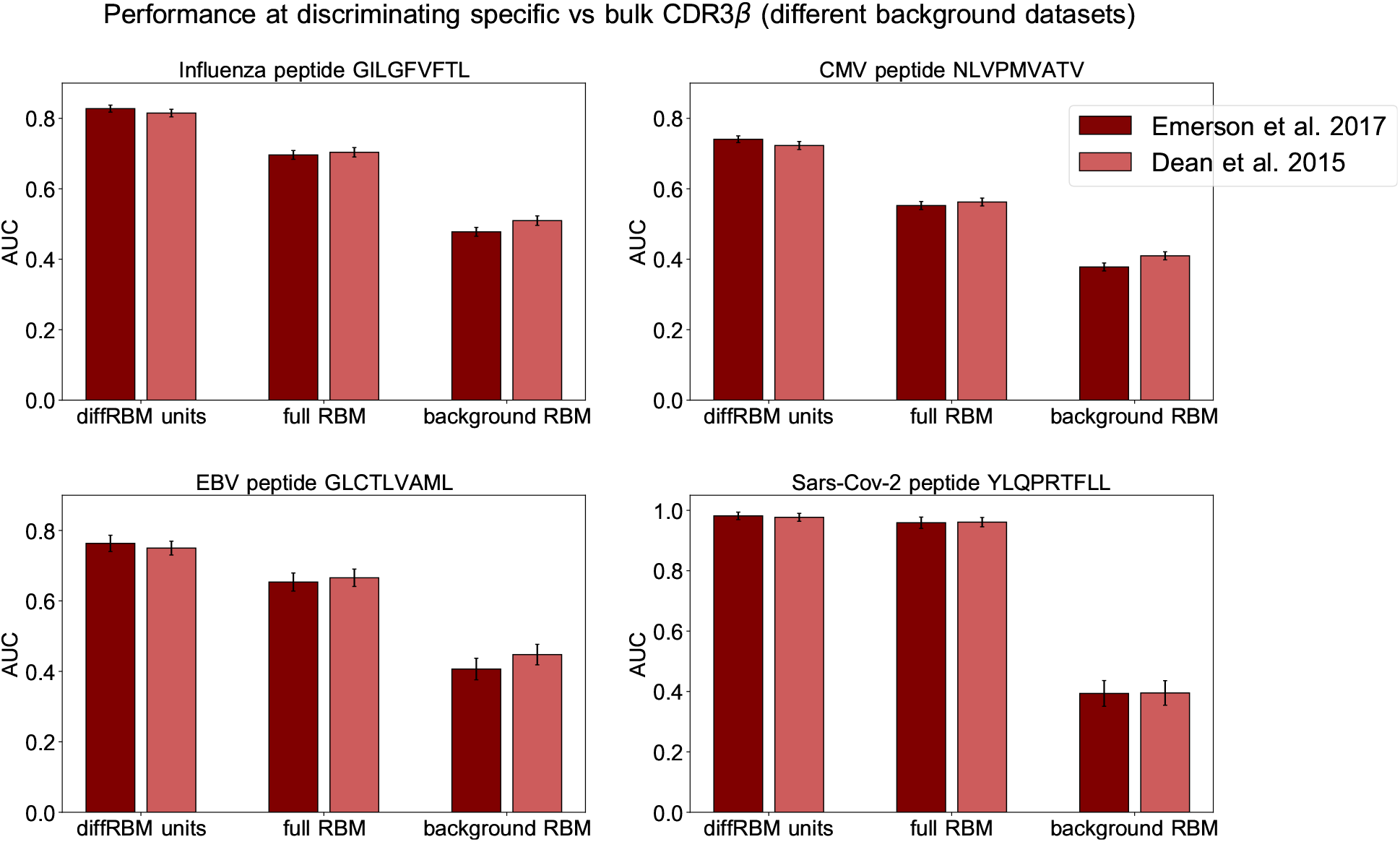
Comparison of performance of differential models of TCR specificity with different background datasets. The plots show, for each epitope, the AUC of discrimination between the epitope-specific and naive sequences for the diffRBM units, full RBM and background RBM in the case where the background dataset is drawn from the [Dean et al., 2015] dataset, compared to the values obtained for a background dataset from [Emerson et al., 2017] (the one used in Figure 6). AUCs are averages over 50 training-test partitions and error bars give the corresponding standard deviation (Methods).

**Figure S18:**
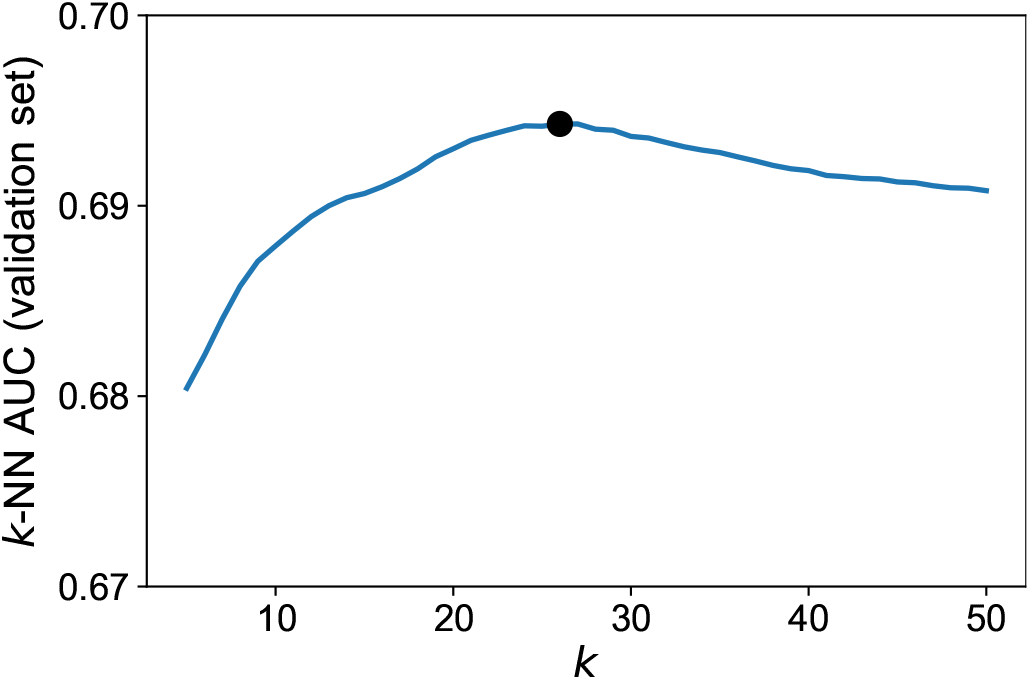
Hyperparametric search of the optimal *k* for the *k*-NN algorithm. The optimal *k* (*k*=26, indicated by the black bold dot) is chosen by looking at the maximal AUC of discrimination between epitope-specific and background sequences in the validation set. For each *k*, the plot shows the average AUC over 50 independent partitions into training and validation sets (Methods). Dataset used: set of sequences specific to the CMV peptide NLVPMVATV 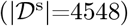.

**Figure S19:**
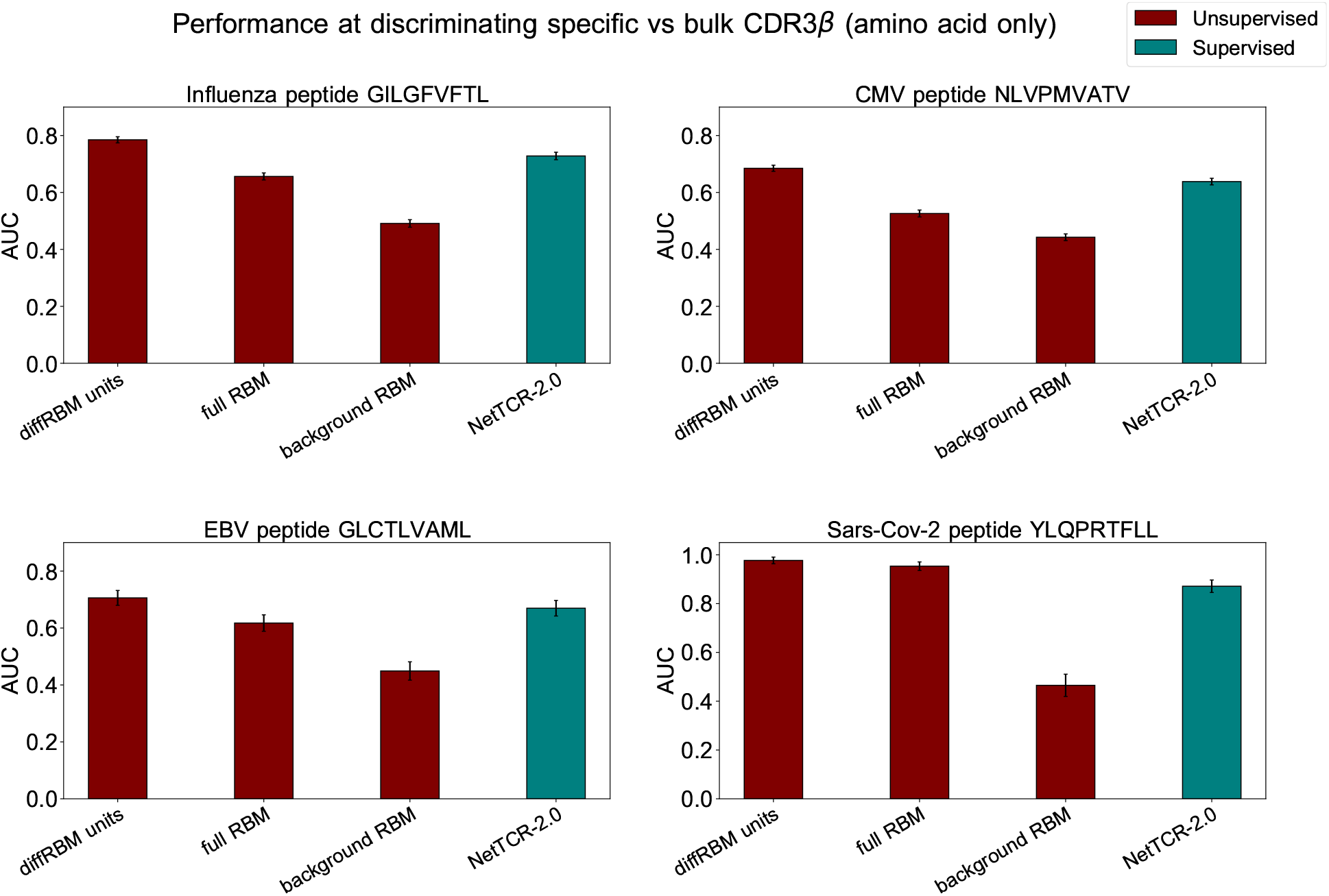
Comparison of performance of models of TCR specificity without V and J type. Given that NetTCR2.0 [Montemurro et al., 2021] does not account for V and J type in its input, we report the performance of the diffRBM units, full RBM and background RBM for the 4 epitope-specific models under consideration when only the CDR3*β* amino acid sequence has been used to train and test the models. AUCs are averages over 50 training-test partitions and error bars give the corresponding standard deviation (Methods).

**Figure S20:**
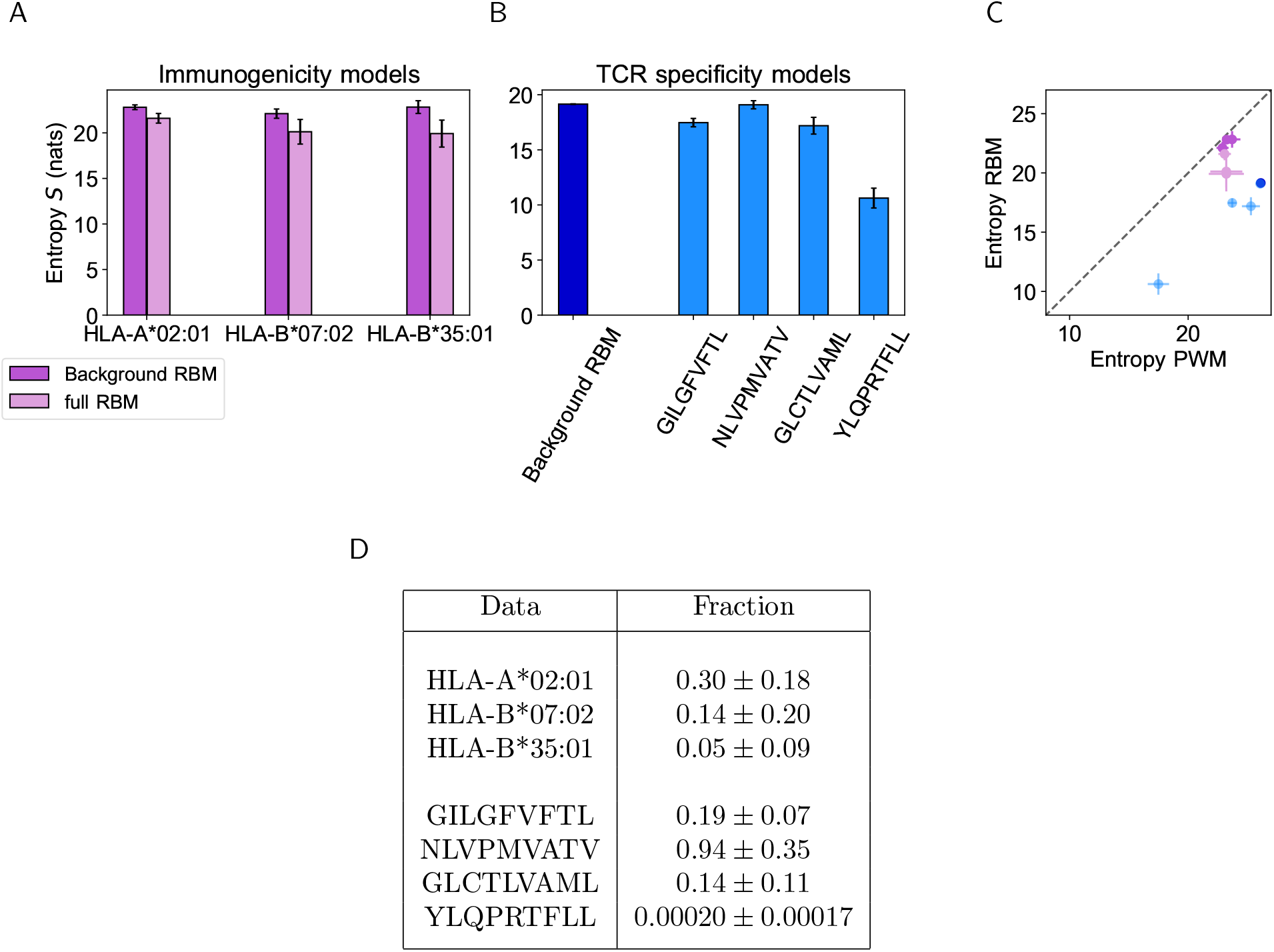
Model-based entropy estimation. A: Entropy *S* (expressed in nats, see Methods) of the space of HLA-specific presented antigens (evaluated by background RBM) and of HLA-specific immunogenic antigens (evaluated by the full RBM) for the 3 HLAs. Error bars represent the uncertainty on the estimated entropy coming from limited sampling and was calculated as in [Marchi et al., 2019]. B: The entropy of the background dataset (CDR3*β* bulk repertoire) obtained from background RBM is compared to the entropy of epitope-specific CDR3*β* obtained from the full RBM models of TCR specificity to GILGFVFTL, NLVPMVATV, GLCTLVAML and YLQPRTFLL. C: The entropies calculated from background RBM and the full RBM plotted in A and B is lower than the one estimated from independent-site models of the same data (Entropy PWM), because RBM models can account for correlations between sequence sites, hence for additional constraints on sequence diversity. Colors are the same as in A, B. D: **Reduction in diversity in selected data.** For each type of data, given the entropy S, exp(S) gives an estimate of the effective size of the corresponding sequence space, hence the ratio exp(*S*^RBM^)/exp(*S*^Background^) gives an estimate of the effective fraction of sequences retained in selected data compared to background ones. The table shows that the reduction in diversity quantified by this fraction is typically less than one order of magnitude, with the exception of the YLQPRTFLL-specific CDR3*β* repertoire. However, the uncertainty from sampling is substantial (especially for the smallest datasets HLA-B*07:02, HLA-B*35:01 and YLQPRTFLL), and prevents us from estimating the reduction in diversity with high accuracy. The first 3 rows refer to the HLA-specific models of immunogenicity (entropy values plotted in A) while the last 4 rows refer to the epitope-specific models of TCR specificity (entropy values in B).

